# *Segatella copri* strains adopt distinct roles within a single individual’s gut

**DOI:** 10.1101/2024.05.20.595015

**Authors:** Xieyue Xiao, Adarsh Singh, Andrea Giometto, Ilana L. Brito

## Abstract

*Segatella copri* is a dominant member of individuals’ gut microbiomes worldwide, especially in non-Western populations. Although metagenomic assembly and genome isolation have shed light on the genetic diversity of *S. copri*, the lack of available isolates from this clade has resulted in a limited understanding of how members’ genetic diversity translates into phenotypic diversity. Within the confines of a single gut microbiome, we have isolated 63 strains from diverse lineages of *S. copri*. We performed comparative analyses that exposed differences in cellular morphologies, preferences in polysaccharide utilization, yield of short-chain fatty acids, and antibiotic resistance across isolates. We further show that exposure to *S. copri* lineages either evokes strong or muted transcriptional responses in human intestinal epithelial cells. Our study exposes large phenotypic differences within related *S. copri* isolates, extending this to host-microbe interactions.

## Introduction

The microbiomes of individuals residing in non-industrialized countries are dominated by a single clade within the Bacteroidota phylum, members of the *Segatella* complex^1,2^. The most abundant and prevalent gut *Segatella* species, *S. copri* (formerly known as *Prevotella copri*^3^), has been associated with various health outcomes, such as glucose intolerance, rheumatoid arthritis and low-grade systemic inflammation with HIV infection^4–8^. However, these associations often conflict, prompting debates regarding the specific role *S. copri* plays within the host intestinal environment. Difficulties in culturing this organism have impeded further exploration into the impact of *S. copri* on the gut ecosystem and host health. It is an obligate anaerobe with understudied nutritional preferences, and, despite the tremendous amount of genetic diversity reported within this clade^1,3,9,10^, until recently, there has been only one strain available from public strain collections. Furthermore, this strain has been recalcitrant to genetic modification and to colonization in mice^11,12^. Overall, these constraints have led to large gaps in experimental evidence supporting these purported roles.

Within the human gut microbiome, each species is often represented by numerous strains exhibiting genomic and functional diversity^13–16^. Lineages of *S. copri* have been identified throughout the world and assembled into metagenomic assembled genomes (MAGs). Using these and available genomes, Blanco-Míguez *et al*. have expanded the *S. copri* complex into 13 distinct species^1,3,17^. Against this backdrop, we asked to what extent the genomic diversity translates into functional and phenotypical variation, relevant both for the bacterium itself and for human health.

In contrast to certain related oral *Segatella* species, *S. copri* poses challenges in cultivation and manipulation. It was not until very recently that people have successfully engineered a small subset of *S. copri* strains. Efforts have also been made to use mouse models to study *S. copri*, but due to limited understanding of its growth preferences and the absence of *S. copri* as a natural member of the mouse gut microbiome, colonization has posed challenges. Gellman *et al*. found that supplementing the mice with plant-derived microbiota-accessible carbohydrates enables colonization and maintenance of *S. copri* strains in mice^12^. In practice, this complicates experimental setups and may not be universally applicable for all research purposes.

Given the challenges of genetic modification and murine colonization, comparative genomics and phenotypic characterization offer an alternative approach to study diverse *S. copri* strains. In this study, we obtained 63 isolates from a fecal sample of Fijian origin. There was considerable genomic diversity among the set of strains inhabiting this single individual. Considering that these strains may co-exist as a result of niche separation, the set of closely related strains provides an interesting vantage point to study the phenotypic diversity across clades. In addition to identifying a new clade of *S. copri*, we find significant differences in their metabolic preferences and production of short-chain fatty acids (SCFAs), in addition to their overall effects on host intestinal cells.

## Results

### Diverse *S. copri* strains isolated from a single individual’s gut microbiome

To explore the diversity of *S. copri*, we employed a refined culturing method on samples enriched in *S. copri*, as determined by metagenomic sequencing. Study participants of the Fiji Microbiome Community Project (FijiCOMP) had high overall burdens of *S. copri* (roughly thirty percent on average)^18^. We selected a 40-year-old female whose microbiome had the highest abundance of *S. copri* (73.9% of the known taxa, as determined by MetaPhlAn2^19^) (**Supplementary Fig. 1A**). By applying our isolation procedures followed by whole genome sequencing, we obtained 63 isolates with high-quality genomes, per field standards^20^, barring 12 isolates that lack full-length 16S rRNA sequences.

The isolates derived from this single individual were remarkably diverse. Despite possessing nearly identical 16S rRNA sequences, the isolate genomes were clustered into six large clades, based to their genomic content and average nucleotide identity (ANI) (**Fig. 1A, Supplementary Fig. 1B**). Phylogenetic analysis yielded the same results (**Supplementary Fig. 1C**). The core genome, defined as those genes represented by over 95% of the isolate genomes, makes up only 423 genes, or 16.8% of each genome on average. Inter-clade ANIs fall below 95%, indicating higher-order relationships beyond the species level (**Supplementary Fig. 1D**). Whereas the isolates obtained from Clade I are likely isogenic (average ANI over 0.999), isolates from Clades III and IV were sparser and showed high intra-clade diversity.

**Fig. 1.**
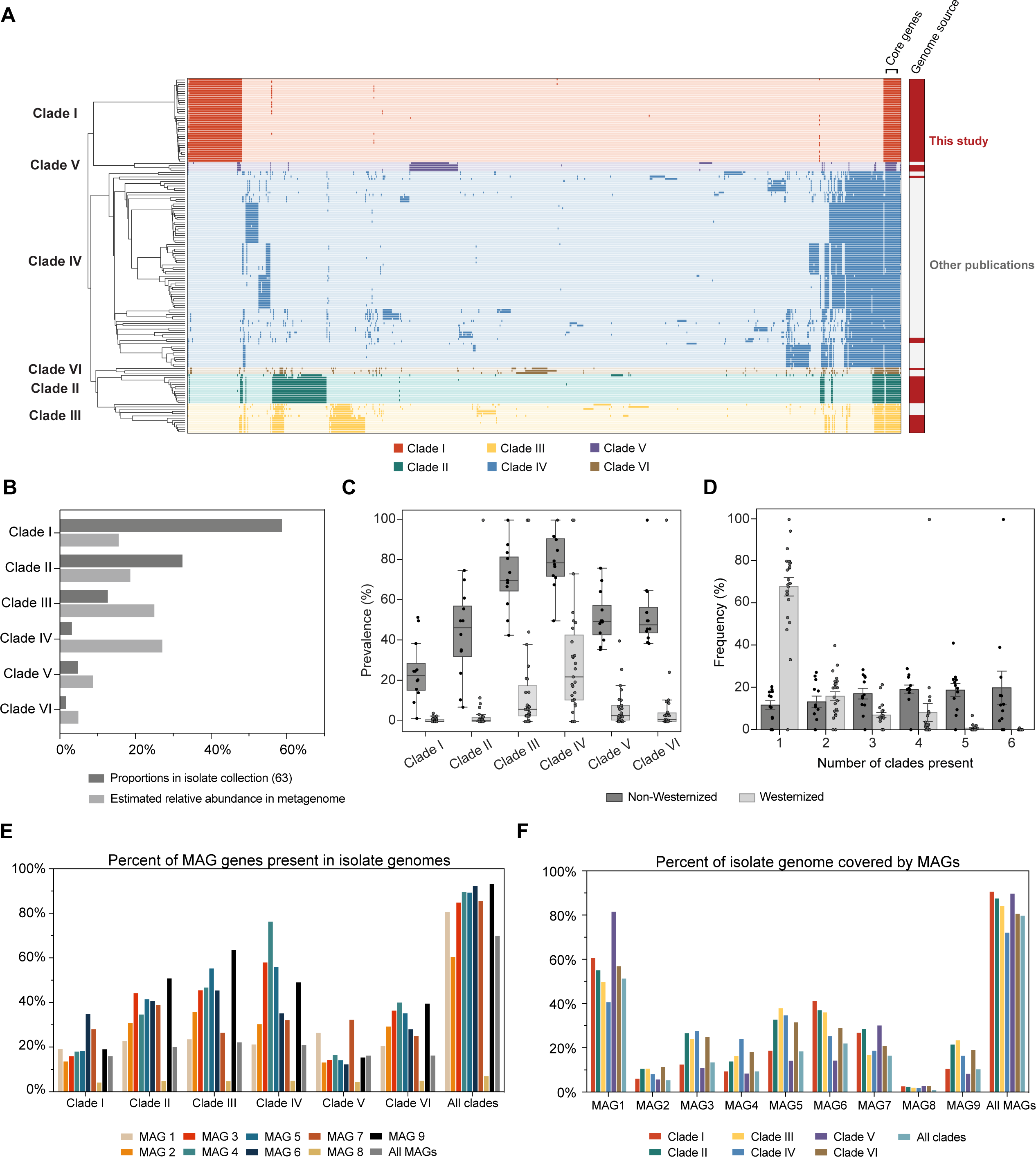
*S. copri* isolates from a single FijiCOMP participant’s gut microbiome cluster into six clades. (A) A cluster map of the genes within the 63 genomes isolated from the Fiji_W2.48.ST FijiCOMP gut microbiome, the genome of *S. copri* DSM18205 downloaded from RefSeq, and the 94 isolate genomes from previous studies. Core genes are defined as those genes appearing in >95% of all isolate genomes. The source of each genome (this study or others’) is indicated in red and gray. The 63 genomes from this study and 94 from others’ studies are indicated in red and gray, respectively. (B) The proportion of isolates from our isolate collection (63 in total) belong to each clade is plotted, in addition to the estimated relative abundance of each clade in the metagenome, as estimated using DiTASiC. (C) Prevalence of *S. copri* clades in fecal metagenomes from Westernized and non-Westernized countries. Each dot denotes the prevalence of the *S. copri* clade in sample from each country. Error bars showing the standard errors. Categorization of the populations in (C) and (D) were provided by curatedMetagenomicData (cMD)^21^. (D) Probability distribution of the numbers of *S. copri* clades present in samples from Westernized or non-Westernized countries. Error bars showing the standard errors. (E) A bar plot showing the percentages of each/all MAG genes identified in each isolate genome according to clade or all clades. (F) A bar plot showing the percentages of genes in each/all clade(s) that can be found in the MAGs.

Furthermore, Clade I isolates constituted an entirely new clade of *S. copri* that did not cluster with any available *S. copri* genomes (**Fig. 1A, Supplementary Fig. 1C**). Interestingly, Clade I isolates were the easiest to isolate, despite their relatively low abundance within the metagenomic sample, 40% of which consists of Clade IV isolates. (**Fig. 1B**). The discrepancy between relative abundance and our ability to culture members of each clade hints at different traits regarding oxygen tolerance and nutritional preferences, resulting in the enrichment of certain clades over the others during isolation.

Given that these strains were derived from a single individual living in the Fiji Islands, we sought to explore how representative these *S. copri* strains were of strains found globally. We examined both core and clade-specific genes in publicly available fecal metagenomic datasets from diverse geographical regions^1,11^. As expected, all six clades have higher prevalence in non-Western countries than Western countries (**Fig. 1C, Supplementary Fig. 2A**). Whereas a substantial number of gut microbiomes from non-Western populations comprise strains from all six clades, the majority of gut microbiomes from Western countries contain only one clade, Clade IV (**Fig. 1D**). The prevalence of most *S. copri* clades decreases gradually with increasing income levels with the notable exceptions of Clade I and V, which have considerable prevalence in some populations with upper middle income (**Supplementary Fig. 2B**). We also checked whether the presence of specific *S. copri* clades were enriched in 17 specific diseases^21^, but no obvious associations were observed (see **Methods**).

### Limitations of metagenomic assembly in distinguishing clades

Metagenomic assembly methods are not often benchmarked against isolate genomes derived from the same microbiome, but rather synthetic microbiomes or assembly statistics. Leveraging the unique opportunity of having both metagenomes and numerous isolates and the extensive use of *S. copri* metagenomic assembled genomes (MAGs), we assessed whether standard pipelines accurately capture genomic content. We applied four established metagenomic assembly pipelines: (1) assembly with MEGAHIT, followed by contig-binning by MetaBAT2^22^; (2) assembly with MEGAHIT, followed by multiple contig binning tools (MetaBAT2, CONCOCT and MaxBin) and bin refinement using DAS Tool^23^; (3) the same as pipeline (2) but using metaSPAdes for assembly^24^; and (4) assembly with metaSPAdes and IDBA-UD, then binning with MetaBAT2^25^. These yielded few if any MAGs (1, 5, 3, and 0, respectively) annotated as *Segatella*, and, despite removing contaminating DNA using MAGpurify^23^, none of these methods produced MAGs meeting field standards (> 90% completeness, < 5% contamination, as determined by CheckM^26^) (**Supplementary Fig. 2C)**. We were not able to determine whether co-assembly produces higher fidelity genomes, yet most of these methods utilize the initial assembly steps above as starting points.

Although the average genome size was similar between the isolate genomes and MAGs, there was greater variability in MAG size and gene content (**Supplementary Fig. 2D**, E), likely indicating incomplete assembly or contamination. The MAGs overall had poor recovery of the *S. copri* genomes: on average, 51.7% of the core genes and 20% of all 11,885 *S. copri* genes were absent from any MAG (**Fig. 1E, F**).

The pangenome analysis incorporating both MAGs and isolate genomes showed that the majority of MAGs did not cluster with the isolate clades (**Supplementary Fig. 2E**). Rather, they formed into a distinct cluster, closer to Clade IV, the most abundant clade in this metagenome. Examination of genes identified in both isolate genomes and MAGs revealed that, on average, the isolate gene pool exhibited higher coverage of MAG genes than vice versa. Particularly, bin 8 displayed notably low coverage for all the isolate clades regardless of its high completeness and low contamination compared to other MAGs (**Fig. 1E, F**). These findings underscore the importance of obtaining whole genome sequencing for identifying novel clades of *S. copri* and obtaining a more complete depiction of the *S. copri* pangenome.

### *S. copri* clades exhibit different cell morphologies

When cultured under the same conditions, the strains exhibit distinct cell morphologies, as observed by scanning electron microscopy, and varied in size (**Fig. 2A, Supplementary Fig. 3A**). While most of the selected isolates grew as rod shapes at stationary phase in Schaedler Broth, substantial filamentation repeatedly occurred in Clade VI (C6-F5) cells (**Fig. 2A, Supplementary Fig. 3B**). Filamentation of bacterial cells can be considered as a stress response and has been reported in several intestinal *Bacteroides* species, albeit not in *S. copri*^27,28^. Filamentation was observed exclusively in Clade VI (C6-F5) cells, but not in other isolates, highlighting another instance of clade-specific responses, even within identical environmental conditions.

**Fig. 2.**
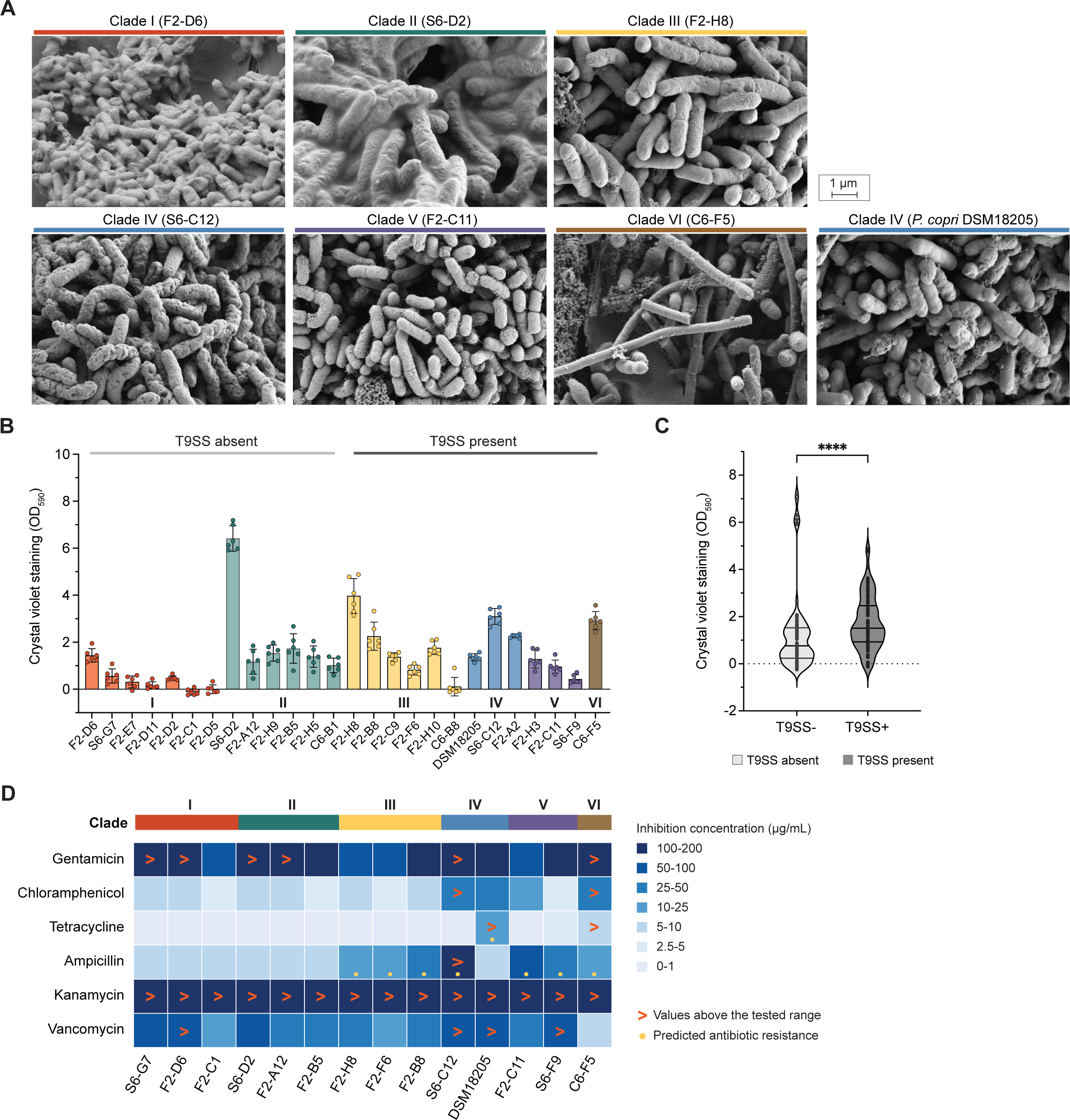
Phenotypic diversity of *S. copri* isolates. (A) Scanning electron microscopy (SEM) images of select *S. copri* isolates, cultured in Schaedler broth until early stationary phase. Isolates names and corresponding clades are labeled on the top. (B) Quantification of biofilm formation by select *S. copri* isolates using crystal violet staining. Optical density at 590nm was measured. Isolates are colored by the clades they belong to (n=6 for each isolate, error bars showing the standard deviations). Dark and light gray bars denote the genomes that with or without T9SS present, respectively. (C) Crystal violet staining readings (OD_590_) of isolates with and without T9SS in the genomes. Statistical significance was calculated by two-tailed Mann-Whitney test (****: p ≤ 0.0001, n=78 for both groups). (D) Minimum Inhibitory Concentration (MIC, μg/mL) ranges of antibiotics on select *S. copri* isolates belong to different clades, as indicated by the color bars. The yellow dots denote the predicted antibiotic resistances as shown in (C). The > symbols indicate that no growth inhibition was observed within the concentration range we tested for the antibiotic.

Surprisingly, Clade II (S6-D2), even after going through a series of fixation and dehydration steps, retained a considerable amount of extracellular substance observable with SEM (**Fig. 2A**). This observation aligns with the findings of a crystal violet assay, which supports the idea that S6-D2 produces significantly higher amounts of biofilm than the other isolates tested (**Fig. 2B**). Biofilm formation often involves self-aggregation in a secreted extracellular polymeric substance that provide metabolic advantages, in addition to antibiotic resistance^29,30^. While oral *Segatella* species have been documented to produce biofilms^31^, such behavior is not observed in gut variants.

The genetic mechanisms regulating biofilm formation in the Bacteroidota phylum, including *Segatella* species, are poorly understood. Type IX secretion systems (T9SS) were previously reported to be involved in biofilm formation of the oral species *Segatella intermedia*^32^. Using TXSscan, we found T9SSs in isolate genomes belonging to Clade III, IV, V, and VI^33^. A significant positive correlation between the presence of T9SS and crystal violet staining was observed, further suggesting involvement of T9SS in regulating biofilm formation in *S. copri* even though it may not be the sole contributing factor (**Fig. 2C**).

### *S. copri* isolates differ in their intrinsic antibiotic susceptibilities

Biofilm formation is often considered one tactic bacteria use to counter the effects of antibiotics. Upon inspection, overall, the *S. copri* genomes we isolated harbor few annotated antibiotic resistance genes. Fourteen of the isolates, the ones belonging to Clade III, IV, V, and VI, harbor the *cfxA6* beta-lactamase gene and the type strain *S. copri* DSM18205 carries a *tetQ* gene providing resistance to tetracycline, both of which were confirmed by higher minimum inhibitory concentrations (MICs) to ampicillin and tetracycline, respectively (**Fig. 2D**, **Supplementary Fig. 3C**). Yet, most of the tested isolates showed low susceptibilities to multiple antibiotics, including gentamicin, chloramphenicol, ampicillin, and vancomycin. In addition, none of the isolates tested was susceptible to kanamycin. Interestingly, a few *S. copri* isolates proved susceptible to gentamicin, which is supposed to be ineffective against anaerobic bacteria due to its oxygen-dependent mechanism of cell membrane penetration^34^. These unexpected results suggest that *S. copri* may possess additional uncharacterized mechanisms of antibiotic resistance. Furthermore, in some cases, isolates from the same clade, even those with high ANI, exhibited different multi-drug resistance profiles, implicating mutations in metabolic genes that can alter antibiotic susceptibilities and/or acquired antimicrobial resistances from the environment or other bacteria species (**Fig. 2D**).

Antimicrobial resistance can be mediated through the acquisition of mobile genetic elements (MGEs). Our isolates harbored various types of MGEs, including transposons, conjugative elements, phage-like elements (lacking phage structural genes), integrons, and mobility islands. Transposable elements were found across all clades, although significantly less in Clade I isolate genomes. Other MGE types were restricted to Clade II, III, IV, and V (**Supplementary Fig. 3D**). MGEs play pivotal roles in bacterial evolution by enabling bacteria to acquire fitness advantages from their environment, although many of the genes within MGEs are poorly annotated. Being equipped with various MGEs likely facilitates the rapid and divergent evolution of *S. copri* clades within the extensive gene pool maintained by intestinal bacteria.

### Clades harbor distinct sets of carbohydrate utilization machineries

As diet has been cited as a main contributor in the colonization of *S. copri*^2,12,35,36^, we directed our focus towards the utilization of dietary fibers as these are a key metabolic feature of members of the Bacteroidota phylum. Polysaccharide utilization loci (PULs), involved in the sensing, transportation, and digestion of available polysaccharides in the environment^37^, have been reported to be differentially distributed among *S. copri* isolates, enabling their digestion of different sets of polysaccharides^9^. We predicted PULs from our isolate genomes by examining loci containing both *susC* and *susD* genes, known to be involved in carbohydrate transfer, and at least one known gene belonging to any carbohydrate active enzyme (CAZyme)^38^ family. The FijiCOMP isolates were rich in PULs, averaging 16 PULs per genome, albeit lower than what was reported in *Bacteroides* species but similar to previous reports in *S. copri*^9,37^ (**Supplementary Fig. 4A**). This amounted to a large number of CAZymes identified within the isolate genomes (**Fig. 3A**), with surprisingly few consistencies across all *S. copri* isolates; only three PUL-associated CAZymes were present in all isolates (GH2, GH3, and GH10). Some PULs are predicted to hydrolyze specific polysaccharides, for instance, the α-mannan-cleaving GH99 exclusively presents in Clade V, which may underlie colonization niches in the gut^39^. Others have enzymes capable of hydrolyzing animal-derived polysaccharides, such as those from sialidase family GH33, which can cleave N-glycolylneuraminic acid (Neu5Gc), a polysaccharide rich in red meat^40,41^.

**Fig. 3.**
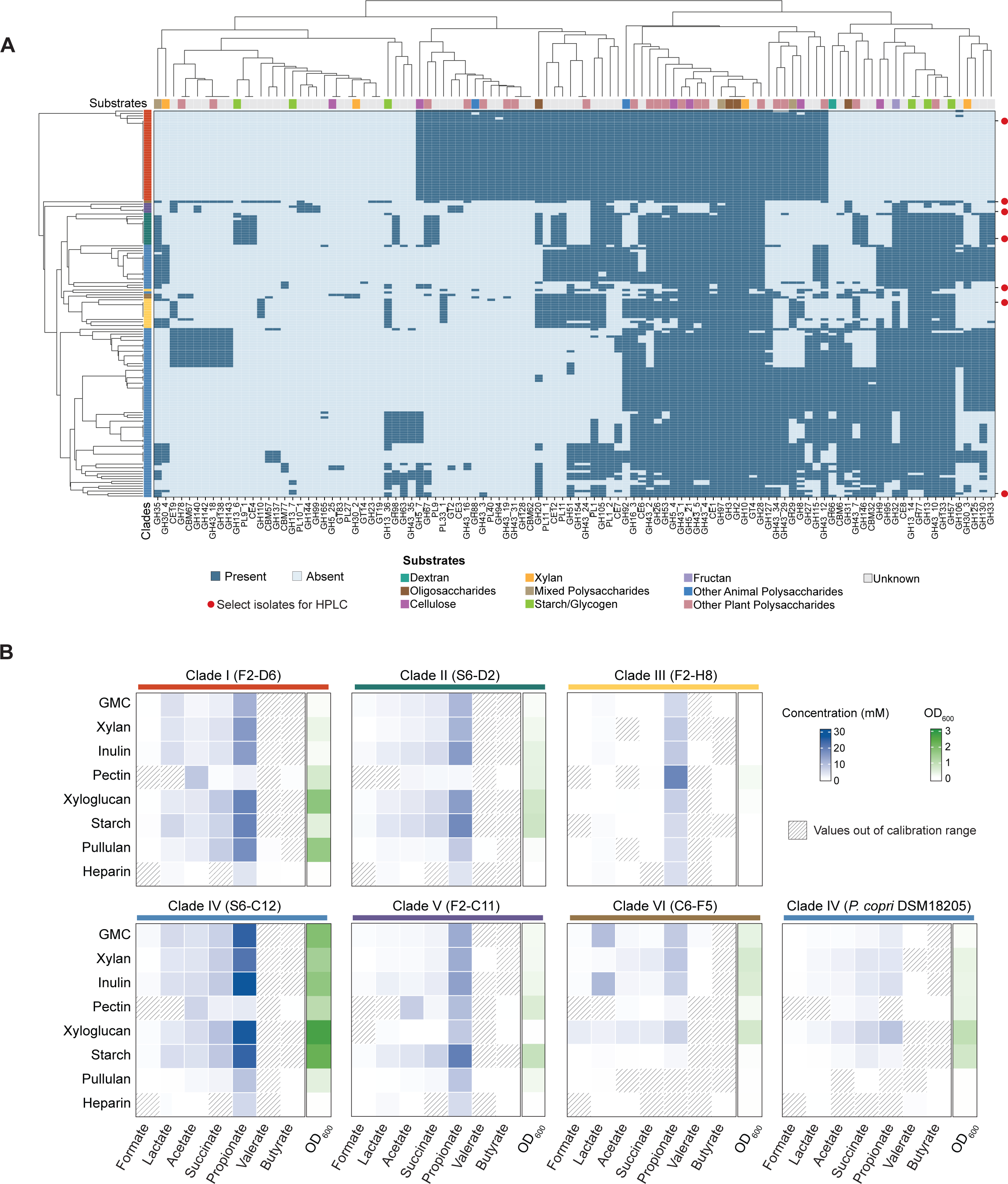
Polysaccharides utilization and acid production by *S. copri* strains. (A) PUL-associated CAZymes identified in *S. copri* isolate genomes. Predicted substrates are depicted at the top. The red dots denote the isolates selected to study the polysaccharides utilization and fermentation products by *S. copri* (from top to bottom: Clade I (F2-D6), Clade VI (C6-F5), Clade V (F2-C11), Clade II (S6-D2), *S. copri* DSM18205, Clade III (F2-H8), and Clade IV (S6-C12)). (B) The growth of select *S. copri* isolates with different carbon sources and the concentration of acids in the spent media at stationary phase measured by HPLC. GMC denote the 1:1:1 mixture of glucose, maltose, and cellobiose, with a same total concentration as the polysaccharides. Side bars showing the growth of bacteria, measured by optical density (OD_600_), using the given polysaccharides or mono-/di-saccharide cocktail as the only carbon source. Shaded cells are values below the calibration range (0-50 mM), indicating possible consumption of acids in the medium.

To test carbohydrate preferences empirically, we performed growth experiments using different polysaccharides as the sole carbon source. We specifically chose six plant-derived polysaccharides, including starch, xylan, and inulin, which are commonly found in the typical Fijian diet, such as taro. Additionally, substrates identified through genomic analysis of PULs, such as xyloglucan, were also included (**Fig. 3B**). To broaden the comparison, we also added a group of mixed mono-/di-saccharides and one animal-derived polysaccharide, heparin. Overall, *S. copri* strains exhibited more robust growth in media containing the plant-derived polysaccharides compared to the two animal-derived polysaccharides, in accordance with previously reported diet correlations^2,36^.

*S. copri* strains showed distinct preferences and abilities in utilizing different polysaccharides (**Fig. 3B**), consistent with the variation observed in PUL content. Five subfamilies of GH5, which is thought to facilitate the degradation of xyloglucan^9,42^, were detected from *S. copri* isolate genomes, whose presence correlated with the growth on xyloglucan (**Fig. 3B, Supplementary Fig. 4B**). Clades achieving moderate to high biomass all had more than three of the five subfamilies including at least one of GH5_2 and GH5_7 subfamilies (**Supplementary Fig. 4B**, C). Multiple subfamilies of CAZyme family GH13, classified as pullulan degrading cazymes^43,44^, were detected from isolate genomes, and, specifically, subfamily GH13_7, annotated as α-amylase, was present in all isolates capable of utilizing pullulan (final OD_600_ >0.1) and was absent from those unable to grow on pullulan. Additionally, Clade IV (S6-C12), which had the highest number of detected CAZymes, was able to grow on all plant-derived polysaccharides, albeit with varying maximum OD_600_. The highest OD_600_, observed with xyloglucan as the carbon source, was 5.5 times higher than the lowest value, observed with pullulan. On the contrary, despite a high number of PULs, Clade III (F2-H8) exhibited mild growth on all of the selected polysaccharides, with the exception of pectin. Surprisingly, none of the strains preferred the mono/disaccharides mixture (GMC) over the plant-derived polysaccharide options, despite cellobiose and glucose being the major carbon sources in M10, the rich medium for the cultivation of *S. copri*.

### *S. copri* isolates produce short-chain fatty acids as a result of polysaccharide degradation

Dietary fibers are degraded by gut bacteria into different types of short-chain fatty acids (SCFAs), which compose up to 10% of the host’s energy budget and confer numerous health benefits, including regulation of host metabolism, immunity, and anti-inflammatory responses^41,45–47^. *Bacteroides* species predominantly produce propionate and acetate^48–50^, whereas intestinal Firmicutes are the main producers of butyrate^51^. Other important fermentation products also include lactate and succinate, the latter of which is reported to benefit the host glucose metabolism by activating intestinal gluconeogenesis^47^. To identify the fermentation products of *S. copri*, we conducted high-performance liquid chromatography (HPLC) analysis on the spent media of seven diverse *S. copri* isolates inoculated with different polysaccharides.

*S. copri* strains showed the ability to produce a variety of SCFAs including formate, lactate, succinate, propionate, and acetate. Butyrate and valerate were also detected from some samples even though at low concentrations. Despite differences in growth of the strains in different carbon sources, strains produced similar ratios of SCFAs (**Fig. 3B**), with some notable differences. Clade I (F2-D6), Clade II (S6-D2), and Clade IV (S6-C12) had similar SCFA profiles. Although Clade III (F2-H8) was unable to grow to a large extent in any of the supplied carbon sources, this isolate was able to produce a considerable amount of propionic acid (18.92 mM), a feat unmatched by the other strains, despite their higher growth. Interestingly, although S6-C12 (Clade IV) and *S. copri* DSM18205 (Clade IV) are genetically very similar (**Fig. 1A**), they exhibited markedly different carbohydrate preferences and SCFA production (**Fig. 3B**). This suggests that the metabolism of SCFAs may be governed by specific sets of enzymes whose presences do not necessarily align with genome clustering.

### Only some *S. copri* isolates induced strong transcriptional responses from intestinal epithelial cells

Given the challenge in colonizing murine models with *S. copri*, especially in light of their highly variable carbohydrate preferences, we chose to analyze host transcriptional responses to *S. copri* using cultured human Caco-2 cells, an intestinal carcinoma cell line. Two-hour incubation with the selected *S. copri* isolates did not induce significant mortality of Caco-2 cells (**Supplementary Fig. 5A**). There were a large number of differentially expressed genes (DEGs) that were strain-specific. However, Caco-2 responses to *S. copri* isolates’ transcriptomes clustered into two groups, with one largely reminiscent of the untreated cells (“hypo-stimulating”), with fewer DEGs compared to the second group (“hyper-stimulating”) (**Fig. 4A, B**). Interestingly, the type strain *S. copri* DSM18205 resulted in few differentially expressed genes (DEGs), despite being reported as pro-inflammatory in previous studies^6,52^.

**Fig. 4.**
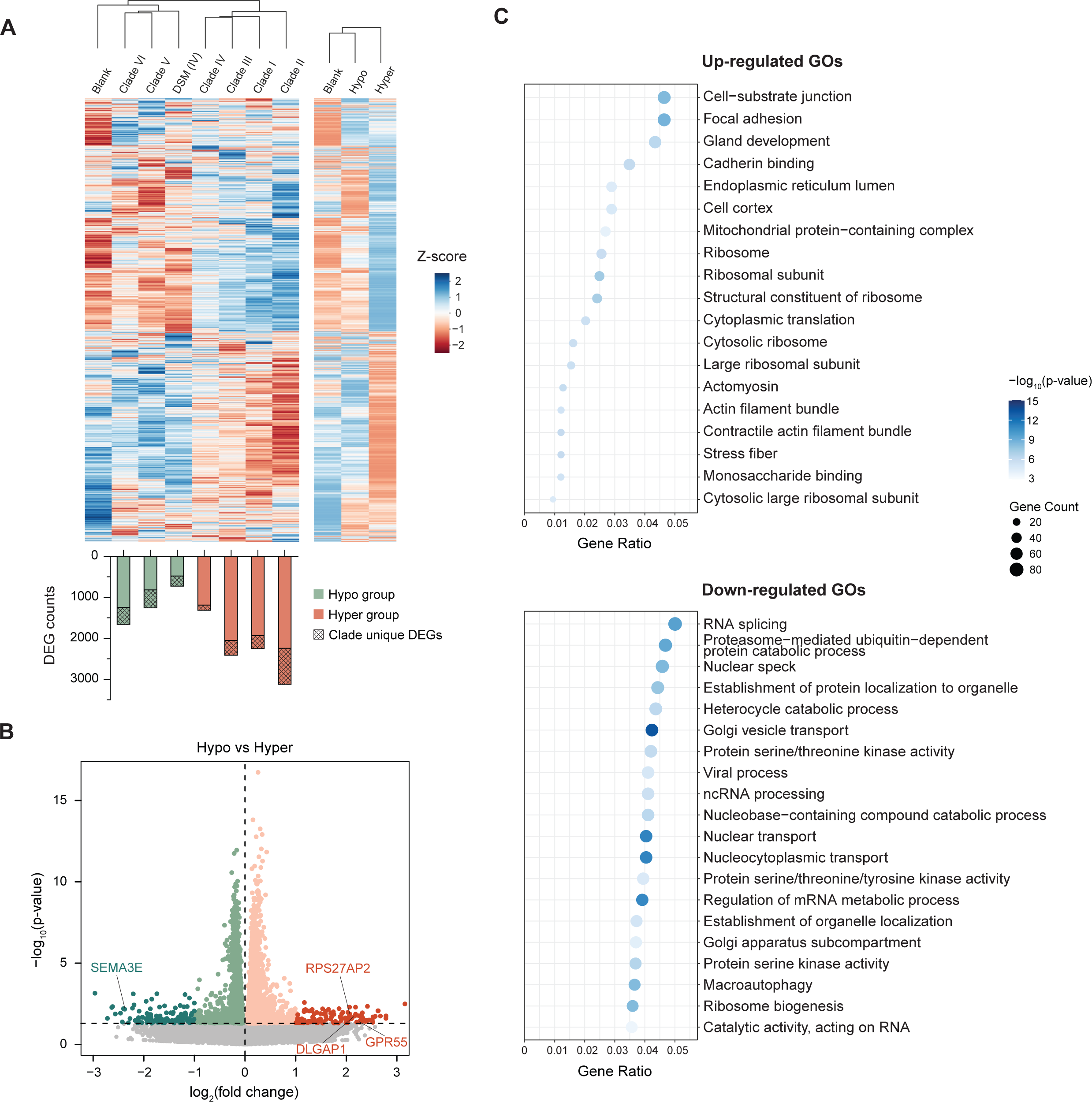
Transcriptomic changes of Caco-2 cells treated with *S. copri* isolates from different clades. (A) The gene expression profiles of Caco-2 cells treated with *S. copri* isolates calculated from the normalized read counts. A Z-score normalization was performed across groups for each gene. Smaller heatmap on the right shows the gene expression files of the Caco-2 cells in Hypo and Hyper groups. The select clade-representative isolates are Clade I (F2-D6), Clade II (S6-D2), Clade III (F2-H8), Clade IV (S6-C12), Clade V (F2-C11), and Clade VI (C6-F5). The bar plot on the bottom shows the number of total and group-unique DEGs found in the indicated treatment conditions compare to the Blank group. (B) Volcano plot visualization the differential expression of genes between Hypo and Hyper groups. Horizontal dash line denoting the p-value cutoff of 0.05. Log_2_(fold change) is used to quantify the differential expression of genes in the Hyper group compared to the Hypo group. (C) The top 20 up-and down-regulated Gene Ontologies (GOs) in Hyper group compared to Hypo group, ranking by p-values. Gene counts are the numbers of DEGs assigned to each GO term.

Among the DEGs with the largest effect sizes between the Hypo and Hyper groups were genes with known association with gastro-intestinal disease. For instance, G protein-coupled receptor 55 (*GPR55*), a gene associated with intestinal inflammation, was found among the DEGs upregulated in the Hyper group^53^. Regulated expression of many long non-coding RNAs (lncRNAs) was also observed including those correlated to gastric cancer and colorectal cancer. For example, *DLGAP1* and *pcsk2-2:1* (or *RPS27AP2*) showed highly increased expression in the Hyper group as well^54,55^. Curiously, among the genes with increased expression in the Hypo group compared to Hyper group (**Fig. 4B**) was *SEMA3E*, a gene whose expression is significantly reduced in ulcerative colitis patients^56^. The regulation of these genes leads to the hypothesis that strains classified into the Hyper and Hypo groups could potentially be disease-promoting or -preventing, respectively. However, further study will be required to understand the full regulatory networks of these genes *in vivo*.

Functional enrichment analyses on the DEGs between the Hyper and Hypo groups identified nearly half of the 19 significantly up-regulated genes in the Hyper group were associated with actin production, implying potential changes in cell morphology, including cell migration or internalization of bacterial

cells. (**Fig. 4C**). Hypothesizing that this may contribute to barrier defects, we tested whether exposure to *S. copri* strains causes hyperpermeability of Caco-2 cell layers. Fluorescein isothiocyanate-labeled dextran (FITC-dextran) of various sizes is used to measure permeability. Yet, no significant changes in cell layer permeability were observed, except for a slight increase in permeability seen for one of the Clade IV isolates (S6-C12) (**Supplementary Fig. 5B)**.

Among all microbiome-derived *S. copri* strains, Clade II and V exhibited the most different gene expression profiles (**Supplementary Fig. 5C**, **Fig. 4A**). Among the 2,972 DEGs identified between Clade II- and Clade V-treated groups, 6 of the top 20 Kyoto Encyclopedia of Genes and Genomes (KEGG) pathways with significantly reduced expression in the Clade II-treated group were associated with immune signaling pathways, including IL-17, NF-kB, Nod-like receptor, C-type lectin receptors, mTOR, and TNF signaling pathway (**Supplementary Fig. 5D**). These findings highlight significant differences of Clade II in modulating the host immune responses. We further annotated the DEGs according to their associations with disease. We observed that, in comparison to the non-treated group, only the treatment with Clade VI bacteria led to elevated expression of numerous genes that are correlated with intestinal/colorectal cancers, amounting to a total of 59 genes (**Supplementary Fig. 6A**).

### Interactions between different isolates led to the changes in relative abundances in co-culture communities

Despite the realization that humans harbor closely related strains with dynamic strain replacement happening occasionally, there is a surprising lack of understanding regarding how closely-related consortia persist^13,57,58^. Within diverse communities, complex competitive interactions are thought to provide stability^59^. Acknowledging the distinct metabolisms and nutritional preferences of different *S. copri* clades, we performed co-culture experiments to decipher pairwise interactions. To identify individual strains, we designed primers to the marker gene, *rplN*, which spanned a region of sufficient genetic diversity to distinguish between isolates. Employing amplicon sequencing, we were able to track the relative abundances of isolates in the co-culture community (**Fig. 5**). In the time frame, most of the pairwise cocultures reached relative abundance ratios different from the starting point (0.5:0.5). Some co-cultures resulted in reduced growth overall, including a strain cocktail involving five strains, suggesting competitive interactions (**Supplementary Fig. 6B**). Clade IV isolate S6-C12 dominated in all combinations, often reaching higher relative abundances than any other co-cultured isolate (**Fig. 5** upper left), whereas Clade V isolate F2-C11 was uncompetitive with all the other strains. These results are in accordance with the estimated clade relative abundances in the fecal metagenomes, of which Clade IV and V presented the highest and lowest abundances, respectively. (Note that Clade VI was not included in the coculture experiment, although its abundance in its source metagenome is below 5%, **Fig. 1B**).

**Fig. 5.**
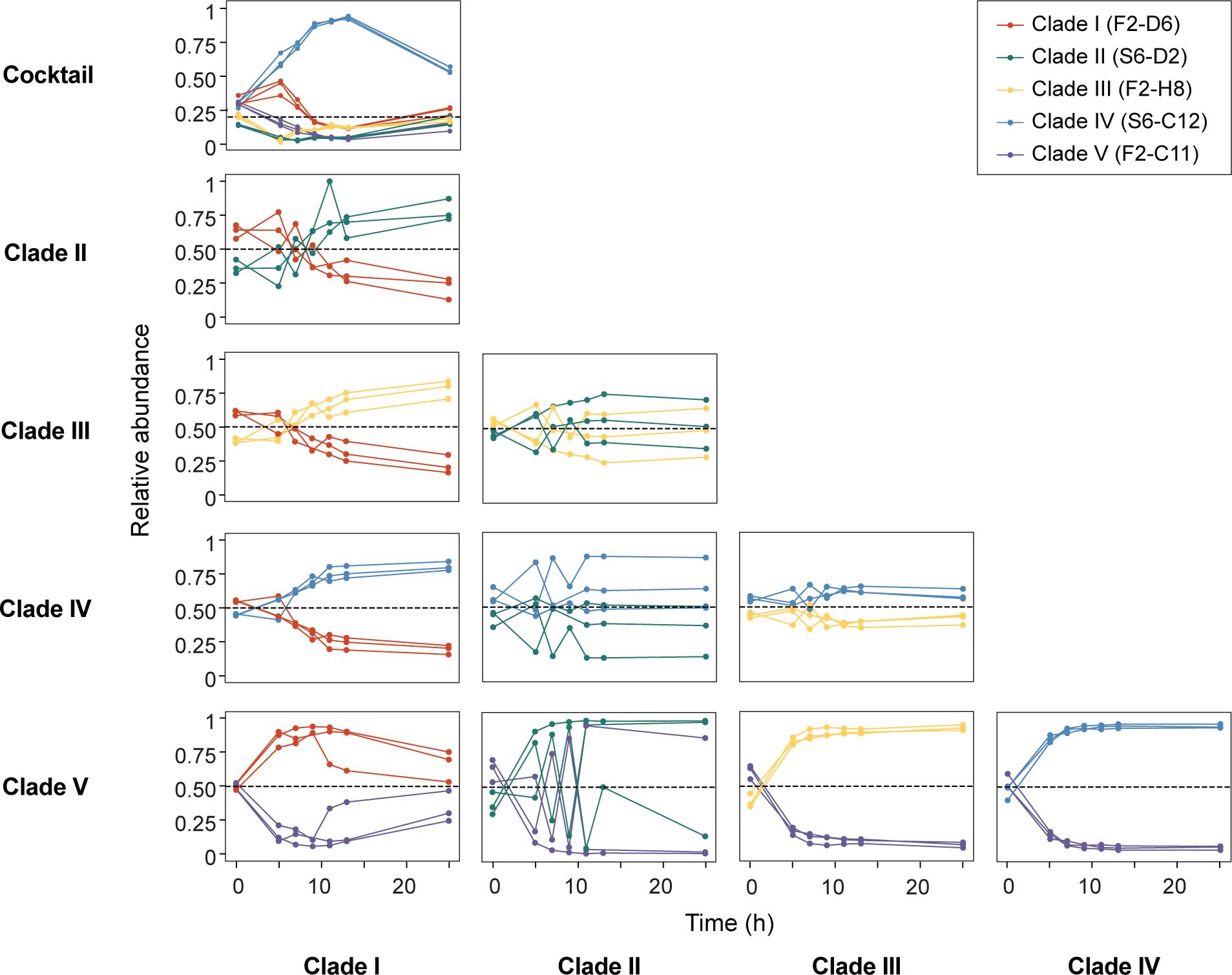
Interaction of *S. copri* strains in coculture communities. (A) Changes in relative abundances of each strain in the cocultures over a 25-hour period. The top plot shows the relative abundances of isolates in the cocktail of the five clade-representing strains, which are Clade I (F2-D6), Clade II (S6-D2), Clade III (F2-H8), Clade IV (S6-C12), Clade V (F2-C11).

We performed a second assay to explore whether any metabolites or proteins were produced by one organism that could promote or inhibit the growth of another, to further probe the interactions between the isolates. *S. copri* isolates cultured in the spent medium of other isolates showed growth inhibition to different extents. The results are consistent with what was observed in the coculture experiment, with Clade IV isolate S6-C12 spent medium showing stronger inhibition of all other strains. Clade I isolate F2-D6 was inhibited by all other isolates (**Supplementary Fig. 6C**). The results indicated that the interactions between cocultured *S. copri* isolates are, at least partially, achieved by secreted small molecule metabolites or proteins during bacteria growth.

## Discussion

*S. copri* was first isolated in 2007, and its genome was made available in 2009^60^. Whereas the study of *Bacteroides* species has been facilitated by the ease at which they can be cultured, since the first *S. copri* isolates were obtained, only one *S. copri* genome has been made available, that from the gut microbiome of a healthy Indian male^61^, and until recently, only one strain was publicly available through commercial strain catalogs. Therefore, most of our knowledge about the role of *S. copri* in the gut microbiome has come from correlative data from case-control metagenomic studies. The intra-species diversity of gut commensal *S. copri* has gained attention in recent years. Tett *et al*. assembled thousands of MAGs from fecal metagenomes in 2019, clustering them into four distinct lineages^1^. Most recently, Blanco-Míguez and colleagues further expanded *S. copri* into 13 distinct species^3^.

We performed whole genome sequencing on 63 of our Fijian *S. copri* isolates clustered into six clades based on their gene content, which revealed the presence a unique clade that was not previously identified. The detected differences in functional gene content are likely reflective of deeper evolutionary relationships rather than recent gene transfer events, as gene clustering largely matches the isolates’ phylogeny. The data presented here, regarding the nutritional preferences, production of metabolites, and interactions with host cells, supports the notion that genomic diversity-driven variations in metabolism and phenotypes provide a possible explanation to the conflicting correlations between *S. copri* and host health. As *S. copri* is being considered to serve as a diagnostic indicator^62,63^ or even a putative therapeutic target^64^, it is critical to understand strain-level effects.

The two prominent genera within the Bacteroidota phylum, *Bacteroides* and *Segatella*, are thought to play a major role in carbohydrate degradation in the gut. *Bacteroides* and *Segatella* species harbor numerous diverse PULs and consistently make up a large portion of individuals’ microbiome composition^65,66^. *Segatella* species are pervasive in developing countries worldwide, where *Bacteroides* dominate in Westernized countries, and it is suspected that diets higher in fiber drive this difference^4,19^. There are some major similarities, including the production of propionate. *Bacteroides-*produced propionate has been reported to play a beneficial role in intestinal immunity and homeostasis^49,50^. Mapping of the central carbon metabolism of *S. copri* DSM18205 (Clade IV) indicated that it is equipped to produce succinate, formate, and acetate with glucose^9,67^. Our results using more complex fibers reveal low production of SCFAs by this strain. On the contrary, our *in vitro* experiments reveal that *S. copri* strains isolated from the Fijian individual broadly produce propionate as a result of degrading plant-derived polysaccharides, among other SCFAs. Our analysis of the PUL-associated CAZymes which have known or predicted substrates, correlates to large extent with the growth observed in each fiber. However, the majority of the CAZymes identified are poorly characterized. Although *S. copri* isolates have universal abilities to produce propionate, preliminary searching of known enzymes responsible of propionate synthesis in *Bacteroides* yielded no homologous genes, suggesting unique, or sufficiently diverged, mechanisms in *S. copri*.

In addition to SCFAs, we previously reported that members of these *S. copri* clades produce diverse and novel sphingolipids, which serve both as important structural components of the cell membrane as well as signaling molecules^10,68^. Due to the structural conservation between bacteria and mammalian sphingolipids, microbiota-produced sphingolipids were reported to be involved in host metabolism and immune homeostasis^68–72^. The production of sphingolipids and SCFAs may underlie some of the transcriptional differences we observed within the *S. copri*-Caco-2 cell coculture experiment presented here. Progress in genetic engineering and/or colonization of *S. copri* in mice models is required to further investigate their roles in host metabolism and immunity. The observed multidrug resistance in *S. copri* clades warrants attention, given the opportunistic pathogenicity of *S. copri*, which has been implicated in bloodstream infections as reported in a prior case study^73^.

Members of the genus *Segatella* have been associated with inflammation in respiratory mucosa and the oral cavity, as well as the vaginal tract^6^. Only recently has their colonization within the gut been associated with gut-associated inflammation and chronic inflammation^6,74^. Given that *S. copri* is also the most abundant gut *Segatella* species^75^, and that *Segatella* are highly prevalent worldwide^1,75^, understanding whether or not all strains of *S. copri* contribute equally to inflammation is of high importance. Confusingly, case-control studies have both implicated and absolved *S. copri* in inflammation-associated metabolic disorders. *S. copri* has been associated with insulin resistance^7^, but not type 2 diabetes^76–78^; with both obesity^79–81^and leanness^82^; and yet is found elevated in patients with rheumatoid arthritis^52^, hypertension^83^, non-alcoholic fatty liver disease (NAFLD)^80,84^ and inflammatory bowel disease^85^. Our study did not find any strong correlation between *S. copri* presence and disease occurrence at the strain-level. A few studies have incorporated mouse models in studying the impacts of *S. copri* on the host, however, those were limited to the type strain *S. copri* DSM18205, which has also been shown to induce pro-inflammatory cytokines IL-6 and IL-23 *in vitro*, thereby promoting Th17-mediated immune responses^6,86^. The colonization of other strains remains challenging as it requires the supplement of plant-derived polysaccharides. We expect that the growing availability of strains and associated genomes will further research into the roles of individual strains in disease and multi-strain consortia that may co-exist in individual’s microbiomes.

## Acknowledgements

This work was funded by grants from the Packard Foundation (to I.L.B.), Pew Charitable Trusts (to I.L.B.) and the National Health Institutes (1DP2HL141007 to I.LB.).

We thank Cornell Center for Materials Research (CCMR) for performing SEM imaging and Weill Cornell Medicine Microscopy and Image Analysis Core Facility for performing TEM imaging. The whole-genome sequencing of *S. copri* isolates was performed by the Biotechnology Resource Center (BRC) at Cornell Institute of Biotechnology.

## Author Contributions

I.L.B., and X.X. contributed to the conceptualization of the work. X.X., A.G., and A.S. built the methodology and performed the research. X.X. and A.S. contributed to the analysis and visualization of the results. X.X. and I.L.B. wrote the original draft and all authors contributed to the review and editing. The work was under the supervision of I.L.B..

## Methods

### Human subject and stool sample collection

Human stool samples used in this study were collected as part of the Fiji Community Microbiome Project (FijiCOMP)^18^. This study was initially approved by the Institutional Review Boards at Columbia University, the Massachusetts Institute of Technology, and the Broad Institute and ethics approvals were received from the Research Ethics Review Committees at the Fiji National University and the Ministry of Health in the Fiji Islands. The Cornell University Institute Review Board additionally approved this study (#1608006528). Human subjects were consented prior to participation in the study. Stool samples were collected into PBS with 20% glycerol within 30 minutes of voiding, preserved in RNALater (QIAGEN), and stored at - 80 °C prior to metagenomic library preparation. The prepared library was sequenced on the Illumina HiSeq2000 platform, 2⨉250 bp paired end reads^18^.

### *S. copri* strains included in this study

The type strain used in this study, *Segatella copri* DSM18205, was purchased from DSMZ. Its genome was downloaded from NCBI RefSeq (GCF_000157935.1). The 63 *S. copri* isolates used in this study were obtained as described in the METHOD DETAILS section. Apart from our own isolate genomes, we included in part of our analysis the isolates from previous publications available at the time this study was conducted. This includes 83 genomes from Tett et al. and 11 genomes from Li et al.^1,11^. The reference genomes of all other *Segatella* species (named as *Prevotella* by NCBI) used in phylogenetic analysis were downloaded from NCBI RefSeq database.

### Bacteria isolation and cultivation

The formula of Modified Medium 10 (M10) agar was modified from a previous study and is prepared as described in **Supplementary file 1**, degassed overnight, and used to isolate *S. copri* from human gut microbiomes ^60,114^. Stool samples were diluted with PBS (10^-1^ to 10^-8^), plated onto M10 agar, and incubated in the anaerobic chamber (3%H_2_, 20% CO_2_, remainder N_2._ Coy Lab Products.) for up to 48 hours. To further identify *S. copri* strains, we performed Polymerase chain reactions (PCR) using previously developed *S. copri* 16S rRNA-specific primers^115^ (primers: g-Prevo-F, g-Prevo-R. **Supplementary Table 1**). Colonies yielded bands with correct sizes were streaked and individual isolates were further verified by Sanger sequencing of the full-length 16S rRNA gene (primers: 27F, 1042R. **Supplementary Table 1**).

When needed, the frozen stocks were inoculated and cultured for 24 hours at 37°C in the anaerobic chamber. In order to get the best revival, the glycerol stocks were first inoculated onto degassed M10 agar plates and then subculture to either BBL™ Schaedler Broth (BD Biosciences or HiMedia) or M10 plates, depending on the requirements of the following experiments. Solid and liquid media were freshly made and stored in the anaerobic chamber overnight to degas before use.

The liquid medium was prepared by dissolving Schaedler Broth powder with water inside anaerobic chamber, adding in 0.05% resazurin, aliquoting desired amount into sample tubes or serum bottles, sealing and autoclaving at 121°C for 15 min. Schaedler Broth from two different manufacturer were used throughout the study as indicated due to the discontinuation of BBL™ Schaedler Broth by BD Biosciences.

### Whole-genome sequencing

*S. copri* DSM18205 and 63 *S. copri* isolates obtained from previous steps were anaerobically cultured on M10 plates for 24 hours and then resuspended in PBS. Genomic DNA was isolated using E.Z.N.A.® Bacterial DNA Kit (Omega). Libraries were prepared using the NEBNext® Ultra™ II DNA Library Prep Kit (Illumina). Libraries were sequenced on the Illumina MiSeq 2⨉250 bp platform.

## Computational methods

### Fecal metagenome processing and analysis

The Fijian fecal sample metagenome was processed using three different pipelines used in previous studies and one pipeline used in the lab to get the MAGs for following analysis.

1. First method was adopted from the study of Pasolli et al.^22^. Contigs shorter than 1000 nucleotides were filtered out after assembling with MEGAHIT. The reads were mapped using bowtie2 (--very-sensitive-local), followed by binning with MetaBAT2 (-m 1500).
2. In the pipeline from *Nayfach et al*^23^, the metagenome was assembled using MEGAHIT followed by contig binning with MaxBin, MetaBAT and CONCOCT. The obtained bins were then refined using DAS Tool and cleaned using MAGpurify.
3. The third method is a pipeline established in the lab combining several previously used and proved pipelines^24^. Briefly, the reads were assembled with metaSPAdes followed by contig binning MaxBin, MetaBAT and CONCOCT binning and bin refinement using DAS Tool.
4. The last method was used by *Chen et al.*^25^. BBTools was used to remove the adapter sequences, contamination from PhiX, and other illumine trace contaminants from the raw reads. Low-quality bases and reads were removed using Sickle. The filtered reads were then assembled using metaSPAdes and IDBA-UD, followed by read mapping using bowtie2 with default parameters. Scaffolds longer than 2.5 kb went through binning by MetaBAT with default parameters. The original pipeline used in Chen et al.’s research includes manual cleaning and curation of the MAGs obtained from steps above, which we skipped in order to compare the performance of pipelines without manual processing.

The qualities of the bins were assessed with and the taxonomic classifications of the MAGs were acquired from CheckM^26^. The composition of the fecal metagenome was profiled by MetaPhlAn2^19^.

### Isolate genome assembly and annotation

The paired-end raw reads were trimmed by Trimmomatic and assembled into genomes using SPAdes v3.10.1^88,89^. Any contigs that are less than 500 bp in length were filtered out. The completeness and quality of assembled genomes were checked with QUAST v4.0 and CheckM v1.0.11 with a contamination cutoff of 5% and completeness cutoff of 95%^26,90^. Open reading frames were predicted by running Prodigal v2.6.3 on obtained genomes^92^. Proteins were annotated from the KEGG (Kyoto Encyclopedia of Genes and Genomes) prokaryotic protein database using DIAMOND v0.9.21 blastx^91^. Sequences were then filtered based on e-value and percent identity. Hits with e-values higher than 1e-5 or less than 30% identity to the reference sequences were removed.

### Phylogenic analysis

*16S rRNA gene:* The 16S rRNA sequences were identified from the genome of each *S. copri* isolate using rnammer v1.2. For the twelve isolates from which we failed to get 16S genes by rnammer, amplification and sequencing of full-length 16S rRNA genes were performed with primers 27F and 1492R (**Supplementary Table 1**) and Phusion® Hot Start Flex DNA Polymerase (NEB). Cycles were performed as: 98°C for 3 min, then 30 cycles of 98°C for 10 s, 60°C for 30 s, and 72°C for 30 s. The PCR products were cleaned using Agencourt AMPure XP Beads (Beckman Coulter) before Sanger sequencing.

The ANI values were calculated using FastANI between each two isolates including the type strain *Segatella copri* DSM18205, of which the genome was downloaded from NCBI RefSeq^97^. Genome based-phylogenomic trees of all *S. copri* isolate genomes and reference genomes of other *Segatella* species were constructed by PhyloPhlAn3 using the default library containing more than 400 marker genes with following options^95^: *--diversity low, --tree raxml*. The tree was annotated and visualized using GraPhlAn^96^.

### Presence of *S. copri* clades in cMD

Genes were called from *S. copri* isolate genomes using prodigal 2.6.3^92^ and blasted against Uniref90 database (September 2023 release) using DIAMOND 2.1.8^91,116^. Genes mapped to the same Uniref90 ID with identity ≥90% were clustered together. If a cluster had genes from ≥95% of genomes of particular clade and has no genes from the other clades, the centroid gene, the gene with longest sequence, was considered as a marker gene for that particular clade^1^. This resulted in 1393 genes for Clade I, 622 genes for Clade II, 437 genes for Clade III, 580 genes for Clade IV, 1206 genes for Clade V, and 490 genes for Clade VI.

cMD^21^ were filtered for fecal metagenomes and only countries with more than 5 samples and samples with ≥20M reads, resulting in 10,400 samples. Read files were adapter trimmed using following parameters with BBTools^109^: *ktrim=r k=23 mink=11 hdist=1 tpe tbo*. To estimate presence of *S. copri* clades in cMD, we mapped cleaned reads to clade marker genes using KMA^113^ and filtered for genes with ≥90% identity and ≥95% coverage. A clade was considered present in the metagenome if ≥ 75% of the clade marker genes were present. The relative abundance of *S. copri* clades in each sample was calculated as:

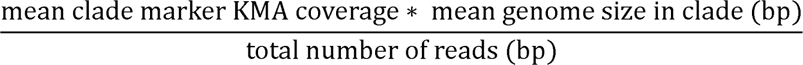

Western and non-western classification was used as provided in cMD and countries were assigned different income-class based on World Bank’s classification downloaded in March 2024 (https://datahelpdesk.worldbank.org/knowledgebase/articles/906519-world-bank-country-and-lending-groups). To assess correlation of disease with prevalence and abundance of *S. copri* clades, we only considered studies with both control and disease samples and performed Fisher’s exact test and Mann-Whitney test respectively.

### Estimation of clade relative abundance

The relative abundance of each *S. copri* clade in the fecal metagenome was estimated using DiTASiC with the default parameters^99^. One isolate genome was picked from each clade as the reference genome. Due to the lower similarity of F2-F6 to other isolates in Clade III, two genomes were picked from this clade as reference genomes (a: C6-B8, b: F2-F6). The proportion of clade in isolate collection was calculated by dividing the total number of isolates by the number of isolates belong to each clade.

### Antibiotic resistance prediction and test

Antimicrobial resistance genes were predicted in the *S. copri* isolate genomes using ABRicate^100^ combining following databases: CARD^117^, EcOH^118^, ARG-ANNOT^119^, Ecoli_VF, VFDB^120^, MEGARES 2.00^121^, Resfinder^122^, PlasmidFinder^123^.

Resistance against various antibiotics were tested using Minimum Inhibitory Concentration (MIC) assays with broth microdilution method^124^. To ensure sufficient growth of *S. copri* isolates, Schaedler broth was used in the MIC tests. Briefly, select *S. copri* isolates were inoculated onto M10 agar plates and cultured for 24 hours. Colonies were collected and resuspended in sterile PBS to 5×10^5^ CFU/mL. Schaedler broth with concentration gradients of select antibiotics were prepared in polystyrene 96 well plates (Costar) and degassed overnight. In the anaerobic chamber, 10 μL of bacteria cell suspension was inoculated into 200 μL of medium per well. Microplates were incubated for 48 hours anaerobically and the optical density at 600 nm was read on Biotek Cytation 5 multimode reader with necessary dilutions. The interpretation of the results referred to the *Reading guide for broth microdilution* and the *Breakpoint tables for interpretation of MICs and zone diameters* and from the European Committee on Antimicrobial Susceptibility Testing (EUCAST)^125^.

### Identification of MGEs from isolate genomes

The MGEs were identified from the isolate genomes following the method provided by by Khedkar et al^126^. Briefly, HMM profiles were built for recombinases using known protein sequences and additional HMMs from Pfam^127^. The recombinases within the isolate genomes were annotated using these HMMs and were mapped to the accessory gene regions to identify recombinase islands. Then the annotated phage structural genes from EggNOG and genes involved in conjugation from TXSscan were mapped to the recombinase islands to assign potential MGEs^33,128^.

### SEM and TEM imaging

For SEM imaging, the bacteria cells were fixed with 2% Glutaraldehyde and 1% OsO4 and transferred to a filter paper after dehydration with serial gradients of ethanol. After overnight critical point drying, the filter papers were sputter-coated with approximately 10nm gold-palladium particles (ratio 60:40) for 60 seconds at 30 mA of current and the images were acquired from the Zeiss Sigma 500 SEM. For *S. copri DSM18205* and isolate F2-D6, the samples were imaged at 0.5kv with a working distance of 2.0 mm and 1.7 mm, respectively. For the rest of the samples, imaging was performed at 1.0 kv with 5.0 mm working distance. Images were acquired with a secondary electron signal using a side angle Everhart-Thornley detector. For TEM imaging, after the same fixation procedures as used for SEM, the pellets were resuspended in 1.5% uranyl acetate and incubated in dark for 1 hour. After dehydration with serial gradients of ethanol, samples were infiltrated and embedded with Quetol 651 for overnight. The samples were viewed on a JEM-1400 transmission electron microscope (JEOL, USA, Inc., Peabody, MA) operated at 100 kV and images were captured on a Veleta 2K x 2K CCD camera (EM-SIS, Germany).

### Biofilm quantification

The biofilm formation by *S. copri* isolates was quantified using crystal violet staining assay^129^. Select *S. copri* isolates were cultured anaerobically in 200 μL of Schaedler Broth on a polystyrene 96-well plate (Costar) for 48 hours. Wells with broth incubated at the same time were used as blanks. The liquid cultures were then aspirated, and plates were dried at 60°C for one hour. Each well was added with 150 μL 0.1% crystal violet solution and stained at room temperature for 15 min. After washing three times with water to remove excess staining, the residual liquid was removed and the plates were dried at 60°C for 10 min. Finally, the biofilm was solubilized and destained with 150 μL 33% acetic acid per well. Absorbance at 590 nm was read on Biotek Cytation 5 multimode reader to quantify biofilm formation. Dilutions were made when necessary.

## Polysaccharides utilization

### CAZyme and PUL predictions

The CAZyme genes in each *S. copri* isolate were predicted and annotated from dbCAN-HMMdb-V11^38^ database using hmmscan (version 3.3) filtered with recommended cutoffs (e-value < 1e-18, coverage > 0.35)^98^. The detected CAZymes were categorized based on the correlated polysaccharide substrates referring to the information in previous studies^130,131^.

### Bacteria cultivation and sample preparation for HPLC

*Segatella* Defined Medium (SDM) was adopted from Defined Minimal Medium Glucose (DMMG) and optimized for the growth of different *S. copri* strains (**Supplementary file 2**)^87,132^. Select *S. copri* strains were first inoculated onto M10 plates and cultured anaerobically at 37°C for 24 hours. Colonies were then collected and resuspended in sterile PBS. After adjusting the OD_600_ to 1.0, 200 uL of the bacteria cell suspension was inoculated into 5 mL SDM with 0.5% (w/v) different carbon sources and cultured anaerobically at 37°C for 48 hours. Blank cultures were inoculated with same amount of PBS. The final optical density at 600nm (OD_600_) of liquid cultures was measured on Biotek Cytation 5 and was calculated by subtracting the blank readings of medium with corresponding carbon sources.

The rest of the liquid cultures was centrifuged at 5000g, 4°C for 15 min to pellet the bacteria cells. The supernatants were collected and filtered through 0.45 um filters. To prepare samples for HPLC, 1 mL of filtered supernatant was transferred to an autosampler vial and then mixed well with 100 uL of concentrated HCl.

### HPLC settings

The HPLC measurement protocol was modified from a previous study ^133^. The Shimadzu HPLC–UV system used consisted of the following modules: a LC-20AD pump, a LC-10AD-VP pump, a DGU-14A degasser, a CBM-20A controlling module, a SIL-20A Auto-sampler, a CTO-20AC oven, an SPD-10A UV detector, and a RF-10A Fluorescence detector. Chromatographic separation was performed using the Hypersil GOLD aQ C18 column (150 mm × 4.6 mm i.d, particle size = 3 µm. ThermoFisher.). The column was thermostatized at 30 °C while running. Two mobile phases were used for the optimal separation of different SCFAs: mobile phase A was 20 mM NaH_2_PO_4_ with pH adjusted to 2.2 using phosphoric acid and filtered with a 0.2 <u>µm</u> filter. Mobile phase B was mobile phase A mixed with acetonitrile (6:4, v/v). The washing buffer was acetonitrile in HPLC water (6:4, v/v). The program of the bi-gradient elution performed is shown in **Supplementary Table 2**. Ten microliters of the prepared samples were injected into HPLC and the UV detector read at a wavelength of 210 nm. The heights of peaks and baselines were acquired from the chromatography and the concentrations of each acid in the samples were calculated from the standard curves described below.

### Calibration and standard curve acquisition

#### HPLC on calibrator solutions

Stock solutions (SS) of select SCFAs were prepared in either HPLC water or 1:1 mix (v/v) of water and HPLC-grade methanol with the concentrations indicated in **Supplementary Table 2** For the acquisition of calibration curves, blank SDM was filter sterilized using a 0.45 <u>µ</u>m filter and was used to prepare calibrator solutions containing following concentrations of all SCFAs: 50 mM, 25 mM, 10 mM, 5 mM, 2.5 mM, 1 mM, 0.5 mM, 0.25 mM, 0.1 mM, 0 mM. Acids with similar elution times were assigned to two different calibration groups (A and B) to better separate the peaks. To run calibrators on HPLC, 1 mL of each calibrator solution was added into an autosampler vial with 100 uL concentrated HCl and vortexed for 15 seconds to fully mix. Ten microliters of the prepared solution were injected and each calibrator solution was run for three times as biological replicates.

#### Calculation of limits of detection and quantification

Linear regression was performed on the peak heights of each SCFA which were acquired as described in the HPLC data analysis section below. The Limit of Detection (LOD) and Limit and Quantification (LOQ) were calculated as suggested by the International Conference on Harmonisation (ICH) as follows where σ is the standard deviation of the response and S is the slope of the calibration curve^134^:

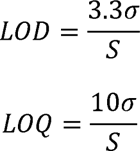

### Bacteria-Human cells co-culture experiment

Co-culture experiment was performed with human Caco-2 cell line at passage 9. Dulbecco’s Modified Eagle Medium (DMEM, Corning) supplemented with 10% Fetal Bovine Serum (FBS) was used as the culture medium. Caco-2 cells (Passage 9) were cultured in 6-well Transwell inserts (Corning, 0.4 uM pore size, Polycarbon membrane) for 4 weeks for differentiation before experiment. On the day of experiment, medium was changed from DMEM supplemented with 10% FBS to pure DMEM.

Seven *S. copri* isolates including the type strain *S. copri* DSM18205 were selected to represent different clades. Bacteria cells were collected from overnight-cultured M10 plates and washed with sterile PBS twice. Bacteria pellets were then resuspended in sterile DMEM medium with OD_600_ adjusted to 1.0. The bacteria suspensions were diluted 1: 50 with sterile DMEM and added into the Transwell inserts at an MOI of 1: 25. Plates were cultured at 37°C aerobically with 5% CO_2_ for two hours. Based on our test, all *S. copri* strains can maintain good viability after two-hour aerobic incubation in DMEM and the viability dropped dramatically afterwards (**Supplementary Fig. 6D**).

### Total RNA extraction and sample quality control

After two-hour incubation, medium was removed from the plates and 1 mL of TRIzol reagent (Invitrogen) was added into each well immediately to collect cells. The total RNAs of Caco-2 cells was extracted by using TRIzol method. Before library preparation, the concentrations of obtained RNA samples were measured using Qubit Fluorometer and the sample integrities and purities were examined using Agilent RNA Sample Quality Control Analysis and agarose gel electrophoresis. Samples with an RNA integrity number (RIN) higher than 8.0 were qualified for library construction.

### Library construction and RNA sequencing

Messenger RNA was purified from total RNA using poly-T oligo-attached magnetic beads. After fragmentation, the first strand cDNA was synthesized using random hexamer primers, followed by the second strand cDNA synthesis using dTTP for non-directional library. After end repair, A-tailing, adapter ligation, size selection, amplification, and purification, libraries were ready for sequencing. The library was checked with Qubit and real-time PCR for quantification and bioanalyzer for size distribution detection. Quantified libraries were pooled and sequenced on Illumina NovaSeq 150bp Paired-end platform, according to effective library concentration and data amount.

### RNA-sequencing Data analysis

Part of the RNA-sequencing data analysis was performed by Novogene.

*<u>Quality control</u>*: The raw data in fastq format was first processed using the fastp software. This step involved extracting clean data (clean reads) by filtering out reads containing adapters, ploy-N sequences, and low-quality reads from the raw dataset. Concurrently, metrics such as Q20, Q30, and GC content were calculated for the clean data. Subsequent analyses were conducted exclusively using this cleaned high-quality dataset.

*<u>Read mapping:</u>* Reference genome and gene model annotation files were downloaded from genome website directly. Index of the reference genome was built and paired-end clean reads were aligned to the reference genome using Hisat2 v2.0.5^135^. Hisat2 was selected as the mapping tool for that it can generate a database of splice junctions based on the gene model annotation file, leading to more accurate mapping results compared to other tools that do not account for splice junctions.

*<u>Gene expression level quantification</u>*: featureCounts v1.5.0-p3 was used to count the numbers of reads mapped to each gene^136^. Then we calculated the Fragments Per Kilobase of transcript per Million mapped reads (FPKM) for each gene based on the gene length and mapped read counts.

*<u>Differential gene expression analysis</u>*: Pairwise differential expression analysis of the treatment conditions (three biological replicates per condition for the eight treatment conditions) was performed using the DESeq2 R package (1.20.0)^101^. The same analysis was also performed between the hyper and hypo group using all samples clustered into each group. The resulting p-values were adjusted using the Benjamini and Hochberg’s approach for controlling the false discovery rate. Genes with an adjusted p-value <=0.05 identified by DESeq2 were assigned as differentially expressed.

*<u>Functional enrichment analysis</u>*: We performed enrichment analyses of Gene Ontology (GO), KEGG pathways, DO (Disease Ontology), and disease related genes with DisGeNET database of differentially expressed genes using the clusterProfiler R package^102^, in which gene length bias was corrected. Functional terms with corrected p-value <=0.05 were considered significantly enriched by differential expressed genes.

### Cell permeability assay

Caco-2 cells (Passage 13) were cultured in 6-well Transwell inserts (Corning, 0.4 uM pore size, Polycarbon membrane) for four weeks to differentiate. The same Bacteria-Human cells co-culture experiment was performed following the protocol above. After two hours incubation, the medium was removed from both the inserts and the basolateral compartments. The cells were washed with PBS and 1.5 mL FITC-Dextran solutions (1 mg/mL) were added to the apical side of the cell layers. The wells were refilled with DMEM and cultured at 37 °C with 5% CO_2_. At 1 h, 2h, and 4 h, 1 mL of samples were collected from the basolateral compartments and refilled with 1 mL fresh DMEM. The concentration of FITC-Dextran was measured on Biotek Cytation 5 (excitation: 490 nm, emission: 520 nm, Bandwidth: 10).

### Primer design and testing for the co-culture interaction experiment

Five isolates from different phylogenetic clades and *S. copri DSM18205* were selected for the co-culture interaction experiment. Marker genes were identified using AMPHORA2 (e-value cutoff = 1e-7) from each isolate genome^93^. A region of *rplN* gene was found to provided sufficient variation to distinguish between each two isolates while a single pair of primers can be used to amplify this region from all select isolates (**Supplementary Table 1**).

To test our method, qPCR was performed on QuantStudio3 Real-Time PCR System using Luna® Universal qPCR Master Mix (NEB) with the following program and different amount of input genomic DNAs were tested: 95°C for 1 min, then 40 cycles of 95°C for 15 s and 60°C for 30 s. The qPCR results confirmed equal amplification of targeting regions from the select isolates, and the differences in input genomic DNAs can be maintained during amplification. Based on the amplification curves, 1 μg of sample DNA was used in library preparation.

### Co-culture interaction experiment

Selected *S. copri* isolates were first inoculated onto M10 plates and cultured anaerobically at 37°C for 24 hours. Colonies were resuspended in sterile PBS and adjusted to have an optical density at 600 nm (OD_600_) of ∼ 1.0. For each pair of organisms, 250 μL suspension of each isolate was mixed and inoculated into 15 mL of warm degassed Schaedler broth. In parallel, 500 μL cultures were set up for each individual organism. Liquid medium inoculated with the same volume of sterile PBS was used as a negative control. For the isolate cocktail group, equal amount of each isolate suspension except the type strain was mix together and 1.5 mL of the cocktail was inoculated into 50 mL freshly made prewarmed Schaedler broth in serum bottle. One mL of liquid culture was drawn at each time point (0h, 5h, 7h, 9h, 11h, 13h, 25h). Cells were collected by centrifuging at 10,000 g for 5 minutes. After removing the supernatant, cell pellets were flash-frozen and stored at -80°C.

### Co-culture interaction library preparation and data processing

*<u>Genomic DNA extraction:</u>* 1 mL of co-culture samples were spun down by centrifuging at 10,000g for 5 minutes, and the cell pellets were used for genomic DNA extraction using Mag-Bind® Bacterial DNA 96 Kit (Omega, Bio-Tek). The DNA extraction was performed following the protocol and automated by epMotion 5075vtc robot (Eppendorf). The concentration of yielded DNAs was measured using Quant-iT™ PicoGreen™ dsDNA Assay Kit (Invitrogen) in 96-well plates.

*<u>Library preparation</u>:* The obtained genomic DNAs were prepared into sequencing libraries and sequenced on Illumina MiSeq 2⨉250 bp platform. The amplicon region of the *rplN* gene was first amplified with a pair of primers containing adapter sequences (rplN-adp-fw, rplN-adp-rev, **Supplementary Table 1**) using the following cycle: 98°C for 30 s, then 22 cycles of 98°C for 10 s, 63°C for 30 s, and 72°C for 10 s, followed by final extension 72°C for 5 min. The PCR products was purified using AMPure XP beads. After clean-up, the DNA concentration was measured using PicoGreen assay as described above, and the yield DNA was diluted to 0.2 ug/ul. In the second PCR, amplicons from different samples were indexed with a set of unique barcodes designed by Diebold et al^137^. One microgram of DNA yielded from the first PCR was input as the template and the cycles were performed as follows: 98°C for 30 min, then 8 cycles of 98°C for 10 s, 64°C for 30 s, and 72°C for 10 s, followed by final extension 72°C for 5 min. Both PCRs used Phusion® Hot Start Flex DNA Polymerase in a 30 μL reaction. Each sample was run as two 15 μL reactions in the second PCR and were pooled together afterwards. The products of the second PCR were again cleaned up by AMPure XP beads purification and the concentrations were determined using PicoGreen assays. Two nanomolar of each sample were pooled together, and the sample purity and fragment size were checked with Fragment Analyzer. The pooled sample was then sequenced on Illumina MiSeq 2⨉250 bp platform. Set-up of PCR reactions, AMPure beads purification, sample dilution and pooling steps were performed on the Eppendorf epMotion 5075vtc robot. PCR reactions were performed on Eppendorf Mastercycler® nexus.

*<u>Cleaning and merging of sequencing reads:</u>* After barcode trimming, the paired-end reads were merged and filtered using USEARCH v11.0.667^94^. The sequencing reads were first merged with the following parameters: length range = expected length of the amplicon ± 20 bp, maximum differences allowed = 22 bases, percent identity ≥ 85%. The merged reads were then passed through the filter with the maximum error threshold of 1.0. Unique sequences were then detected from the filtered reads and the count of each unique sequence was calculated.

*<u>Relative abundances calculation</u>:* The unique sequences from each sample were aligned to the *rplN* genes extracted from isolate genomes to determine the corresponding source strain of each unique sequence. The percentage of unaligned reads or reads from unexpected source strain was calculated to test the quality and purity of the samples. Three samples were found with either inadequate merged reads or overabundant reads from unexpected source strains and therefore were removed from the following analyses. Then the counts of the unique sequences provided by USEARCH were used to calculate the relative abundances of the two strains in each co-culture sample.

### Isolate supernatants inhibitory Assay

*<u>Supernatant collection</u>:* To collect culture supernatants, *S. copri* DSM18205 strain and eight select isolates representing different genomic groups were first inoculated onto fresh-made M10 plates and cultured anaerobically at 37°C for 24 hours. Colonies were collected from the plates and resuspended in sterile PBS with OD_600_ readings adjusted to ∼ 1.0. Three milliliter of cell suspension of each strain was inoculated into 100 mL warm degassed Schaedler broth in a serum bottle and anaerobically cultured for 18 hours until they have reached the early stationary phase. Medium inoculated with sterile PBS was processed and used as the negative control in the experiments. The OD_600_ readings of samples at the time of collection were measured using a SpectraMax M3 microplate reader. Liquid cultures were centrifuged at 7000g for 10 minutes to spin down the bacteria cells. The supernatants were collected and passed through 0.22 μm filter cups to sterilize. The supernatant filtrates were flash-frozen and stored at -80°C for future use.

*<u>Inhibitory assay</u>:* Before inoculation of the *S. copri* strains, 5 mL of each sterile supernatant was combined with 5 mL of autoclaved Schaedler broth in a sample tube and degassed in the anaerobic chamber overnight. The nine select strains were streaked onto fresh-made M10 plates and cultured anaerobically at 37°C for 24 hours. Colonies were collected from the plates and resuspended in sterile PBS with OD_600_ readings adjusted to ∼ 1.0. Three hundred microliters of cells suspension of each strain was inoculated into culture tubes containing different supernatants and fresh medium and cultured anaerobically at 37°C. The OD_600_ values were measured 24 hours after inoculation using the SpectraMax M3 microplate reader.

### Statistical analysis

Statistical analyses of Fig. 2C and 5C were performed in GraphPad Prism. The differences of two groups were compared using two-tailed Mann-Whitney test and multiple t-test with Bonferroni correction, respectively. The methods, sample sizes, and significance levels are indicated in the figure legends.

## Materials Availability

Fecal isolates of *Segatella copri* acquired in this study can be provided upon request.

## Data Availability

The Whole-genome sequencing data of *Segatella copri* isolates passed filtering were deposited to NCBI BioProject PRJNA217052. The metagenome data of Fijian sample Fiji_W2.48.ST was obtained in a previous work and is available under the same BioProject. The accession numbers of all corresponding SRAs are listed in **Supplementary Table 3**.

## Correspondence

Further information and requests for resources and reagents should be directed to and will be fulfilled by the Lead Contact, Ilana L. Brito.

