## Supplemental files and figures for "*Segatella copri* strains adopt distinct roles within a single individual’s gut"

##### Supplementary figure legends

**Supplementary Fig. 1 | Composition of the fecal metagenome and phylogenetic analysis of *S. copri* isolates**

(A) The genus level microbiome composition of the Fijian fecal sample used for *S. copri* isolation in this study. The relative abundance was calculated by MetaPhlAn2. *Segatella* genus comprises 73.9% of known taxa or 23.0% in total. The third largest portion comprises an unclassified genus of Lachnospiraceae family.
(B) Average nucleotide identities (ANIs) between *S. copri* genomes calculated by FastANI. The minimal ANI value was used as the cutoff of the color bar on the heatmap.
(C) The phylogenetic tree of *S. copri* isolates from both this study and previous studies constructed based on the marker genes by PhyloPhlan3.
(D) The mean values of ANIs between genomes within the same (intra-) or different (inter-) clades. Error bars showing standard deviations.

**Supplementary Fig. 2 | Prevalence of *S. copri* clades in different populations and comparison with metagenome assembled genomes (MAGs)**

(A) Presence of the *S. copri* clades in the cMD fecal metagenomes from different countries. The origins of the fecal metagenomes and income levels corresponding to (C) are indicated by the side bars[21]. Star denote the unique clade identified by this study.
(B) Prevalence of *S. copri* clades in samples from countries categorized by income levels, designated by the World Bank's classification. Error bars showing the standard errors.
(C) The completeness and contamination of MAGs acquired from the Fijian fecal metagenome, calculated by CheckM.
(D) The sizes of the *S. copri* isolate genomes, including the type strain genome from RefSeq, and the MAGs acquired in this study. Error bars showing the standard deviations (n=64 for the isolate genomes, n=9 for the MAGs).
(E) Clustering of MAGs together with the *S. copri* isolate genomes based on their gene content.

**Supplementary Fig. 3 | Differences in cell morphology and functional genes among *S. copri* isolates.**

(A) The cell lengths of different *S. copri* strains measured from the SEM images, n=13 for each strain.
(B) Transmission electron microscopy (TEM) images of Clade VI (C6-F5) bacteria cells
(C) Predicted antimicrobial resistance genes in *S. copri* isolate genomes using ABRicate.
(D) Presence of MGEs in *S. copri* isolate genomes. Numbers on the color scale denote log(1+n) where n is the number of corresponding MGE categories contained in the genome. IS/Tn: Insertion sequence elements and Transposons.

**Supplementary Fig. 4 | Presence of PULs and CAZymes in *S. copri* isolate genomes**

(A) Number of predicted PULs in each isolate genome.
(B) Presence of all identified CAZymes in *S. copri* isolate genomes.
(C) Presence (blue) of GH5 subfamilies in the *S. copri* strains used in the polysaccharide utilization experiment. Row on the top shows the growth ($OD_{600}$) of each isolate using xyloglucan as the only carbon source.

**Supplementary Fig. 5 | Transcriptomic changes of Caco-2 cells induced by select *S. copri* isolates.**

(A) Viability of Caco-2 cells after 2-hours coculture with select *S. copri* strains measured by trypan blue staining and read by Invitrogen Countess II Automated Cell Counter (error bars showing the mean ± SD (n=3)).
(B) Results of Caco-2 cell permeability assay with FITC-Dextran of different sizes (FD4k: 4 kDa, FD40k: 40 kDa, FD70k: 70 kDa). Fluorescence intensity of the medium in the basolateral compartments of the Transwell plates were read 4 hours after the stimulation of *S. copri* strains. Statistical significance was calculated by multiple t-test with Bonferroni correction (**: $p \leq 0.01$. Error bars indicating mean ± SD (n=3)).
(C) Volcano plot visualization the differential expression of genes between Clade II (S6-D2) and Clade V (F2-C11) treatment groups. $Log_2$(fold change) is used to quantify the differential expression of genes in Clade II (S6-D2) group compared to Clade V (F2-C11) group.
(D) The top 20 up- and down-regulated KEGG pathways in Clade II group comparing to Clade V group, ranked by gene ratios. Gene counts are the numbers of DEGs assigned to each pathway.

**Supplementary Fig. 6 | Interactions in the *S. copri* isolate cocultures.**

(A) The top 20 up- and down-regulated Disease Ontologies (DOs) in treatment group of Clade VI (C6-F5) compared to the Blank group, ranking by p-values. Gene counts are the number of DEGs assigned to each DO.
(B) The final biomass of the liquid cultures in the coculture interaction experiment quantified by $OD_{600}$. Orange rectangles indicate cultures of single isolates, cocktail: cocktail culture of five clade-representing *S. copri* isolates. Other cells are pairwise co-cultures of the two corresponding stains (n=3 for each culture).
(C) Heatmap showing the growth of *S. copri* isolates ($OD_{600}$) on 1:1 mixture of fresh Schaedler broth and spent medium collected at early stationary phase of other isolates. For the PBS group, a 1:1 mixture of fresh Schaedler broth and PBS solution was used. The fresh medium group used 1X fresh Schaedler broth.
(D) Viability of *S. copri* isolates in DMEM during aerobic incubations mimicking the Caco-2 cell coculture experiment settings. Cell suspensions were plated on M10 agar with dilutions and Colony forming units (CFUs) were calculated (n=3).

#### SUPPLEMENTAL DATA

**Supplementary file 1. Medium 10 (M10) formula and preparation**

| Components | Amount in 1L medium |
|---|---|
| Water | 960 mL |

| | |
|---|---|
| Glucose | 0.5 g |
| Cellobiose | 0.5 g |
| Soluble starch | 0.5 g |
| Minerals stock | 20 mL |
| L-cysteine-HCl ($H_2O$) | 0.5 g |
| Resazurin | 2 mL |
| $Na_2CO_3$ | 4 g |
| Trypticase peptone | 2 g |
| Yeast extract | 0.5 g |
| Volatile fatty acid mix | 3.1 mL |
| Hemin | 20 mL |
| Agar | 20 g |
| $K_2HPO_4$ (after autoclave) | 1 mL |

**Preparation**
Dissolve all the components except $K_2HPO_4$ in 960 mL distilled water and autoclave at 121°C for 15 min. Cool the medium down to 50-60°C, add in 1 mL $K_2HPO_4$ and mix well. Pour the medium into plates. After the plates are solidified, transfer into anaerobic chamber to degas overnight.

**Stock solutions**

Minerals solution (1L):

Except $K_2HPO_4$ which is added separately after autoclave, the minerals are prepared into a stock solution and store at room temperature. Mix well before use.

| | Amount in 1L solution (g) | Final concentration in stock (M) |
|---|---|---|
| $KH_2PO_4$ | 0.177 | $1.3\times10^{-3}$ |
| NaCl | 0.044 | $7.6\times10^{-4}$ |
| $(NH)_2SO_4$ | 0.449 | $3.4\times10^{-3}$ |
| $K_2HPO_4$ (separately) | 0.296 | $1.7\times10^{-3}$ |
| $CaCl_2$ | 0.046 | $4.1\times10^{-4}$ |

| | | |
|---|---|---|
| $MgSO_4 \bullet 7H_2O$ | 0.094 | $3.8\times10^{-4}$ |

Fatty acids mix:

| | Volume in the stock (37 mL) |
|---|---|
| Acetic acid | 17 mL |
| Propionic acid | 6 mL |
| Butyric acid | 4 mL |
| Isobutyric acid | 1 mL |
| n-valeric acid | 1 mL |
| Isovaleric acid | 1 mL |
| DL-alpha-methylbutyric acid | 1 mL |

Hemin stock solution (0.5 mg/mL):
Dissolve 50 mg hemin in 1 ml 1 N NaOH; make up to 100 ml with distilled water. Store refrigerated and avoid light.
$K_2HPO_4$ stock solution (296.106 g/L):
Dissolve 14.8053 g $K_2HPO_4$ in 50 ml distilled water, filter through 0.22 μm filter tube (Steriflip, Millipore). Add 1 mL per 1 L medium after autoclave.
Resazurin stock solution (0.05%, nonsterile):
Dissolve 25 mg resazurin in 50 ml distilled water.

**Supplementary file 2. *Segatella* Defined Medium (SDM) formula and preparation**

| | Per Liter |
|---|---|
| *Carbon source | 5g |
| $(NH_4)_2SO_4$ | 1 g |
| $KH_2PO_4$ | 0.9 g |
| NaCl | 0.9 g |
| Cysteine-HCl x $H_2O$ | 0.5 g |
| SCFAs | 183 uL |
| Hemin | 20 mL |
| $CaCl_2$ | 1 mL |
| Chlorides solution | 1 mL |
| Resazurin | 2 mL |
| *Vitamin K1 | 200 uL |
| Wolfe’s vitamin mix | 1 mL |

| | |
|---|---|
| Water | 925 mL |

*Filter-sterilize stock solution and add after autoclave.

**Preparation**

Dissolve all the components in 925 mL distilled water, adjust the pH to 7.2 using 50% $Na_2CO_3$. After autoclaving at 121°C for 15 min, cool down the medium and add in vitamin K1 and carbon source solutions as described in the formula.

**Stock solutions:**

*Chlorides (nonsterile)*

| | Gram per liter |
|---|---|
| $CaCl_2$ | 20 |
| $MgCl_2$ x $6H_2O$ | 20 |
| $MnCl_2$ x $4H_2O$ | 10 |
| $CoCl_2$ x $6H_2O$ | 1 |

*$FeSO_4$ (nonsterile)*

Make the stock solution by dissolving 4 g of $FeSO_4$ x $7H_2O$ into 1 L of MilliQ water. Mix before use.

*Vitamin K1*

Dissolve 0.1 ml of vitamin K1 in 20 ml 95% ethanol and filter sterilize. Store refrigerated avoid light.

*SCFAs and Hemin*

Both SCFA and hemin solution have the same composition as used in M10 agar (**Supplement file 1**).

**Supplementary Table 1. Primers used in this study**

| Name | Sequence | Usage |
|---|---|---|
| rplN_adp_fw | ACACGACGCTCTTCCGATCTGTGGTACAGGCCGTCGTTAC | Co-culture interaction library preparation (Adapters underlined) |
| rplN_adp_rev | GAGTTCAGACGTGTGCTCTTCCGATCTTCAGGTGCCAAAGAAACGAC | |
| g-Prevo-F | CACRGTAAACGATGGATGCC | *Segatella*-specific primers |
| g-Prevo-R | GGTCGGGTTGCAGACC | |
| 27F | AGAGTTTGATCCTGGCTCAG | Full-length 16S rRNA gene |
| 1492R | GGTTACCTTGTTACGACTT | |

**Supplementary Table 2. HPLC bi-gradient elution program for the analysis of SCFAs**

| Time (min) | Mobile phase A (%) | Mobile phase B (%) | Flow rate (mL/min) |
|---|---|---|---|
| **0** | 100 | 0 | 0.6 |
| **6** | 100 | 0 | 0.6 |

| | | | |
|---|---|---|---|
| **6.5** | 75 | 25 | 0.6 |
| **8.5** | 75 | 25 | 0.6 |
| **9** | 75 | 25 | 1.25 |
| **10.5** | 75 | 25 | 1.25 |
| **11** | 50 | 50 | 1.25 |
| **16** | 50 | 50 | 1.25 |
| **16.5** | 100 | 0 | 0.6 |
| **17** | 100 | 0 | Stop |

**Supplementary Table 3. Sequencing data accessions**

| **Genomes/metagenome** | **Accession** |
|---|---|
| Fiji_W2.48.ST | SRS475594 |
| *S. copri* DSM18205 | GCF_000157935.1 |
| S6-C1 | SRR26987610 |
| S6-C2 | SRR26987609 |
| S6-C3 | SRR26987598 |
| S6-C8 | SRR26987587 |
| S6-C12 | SRR26987576 |
| S6-D1 | SRR26987565 |
| S6-D2 | SRR26987554 |
| S6-D7 | SRR26987550 |
| S6-D10 | SRR26987549 |
| S6-E7 | SRR26987548 |
| S6-E10 | SRR26987608 |
| S6-E12 | SRR26987607 |
| S6-F1 | SRR26987606 |
| S6-F9 | SRR26987605 |
| S6-F11 | SRR26987604 |
| S6-G6 | SRR26987603 |
| S6-G7 | SRR26987602 |
| S6-G8 | SRR26987601 |
| S6-H5 | SRR26987600 |
| C6-A1 | SRR26987599 |
| C6-A2 | SRR26987597 |
| C6-B1 | SRR26987596 |
| C6-B3 | SRR26987595 |
| C6-B8 | SRR26987594 |
| C6-D1 | SRR26987593 |
| C6-D3 | SRR26987592 |
| C6-F5 | SRR26987591 |
| C6-H5 | SRR26987590 |
| C6-H7 | SRR26987589 |
| F2-B8 | SRR26987588 |
| F2-A2 | SRR26987586 |

| | |
|---|---|
| F2-C5 | SRR26987585 |
| F2-H6 | SRR26987584 |
| F2-C12 | SRR26987583 |
| F2-A9 | SRR26987582 |
| F2-C6 | SRR26987581 |
| F2-E7 | SRR26987580 |
| F2-H7 | SRR26987579 |
| F2-D11 | SRR26987578 |
| F2-A11 | SRR26987577 |
| F2-C9 | SRR26987575 |
| F2-E11 | SRR26987574 |
| F2-H9 | SRR26987573 |
| F2-F2 | SRR26987572 |
| F2-A12 | SRR26987571 |
| F2-C10 | SRR26987570 |
| F2-F5 | SRR26987569 |
| F2-H10 | SRR26987568 |
| F2-F6 | SRR26987567 |
| F2-B4 | SRR26987566 |
| F2-C11 | SRR26987564 |
| F2-H1 | SRR26987563 |
| F2-H2 | SRR26987562 |
| F2-B5 | SRR26987561 |
| F2-D2 | SRR26987560 |
| F2-H11 | SRR26987559 |
| F2-B12 | SRR26987558 |
| F2-D5 | SRR26987557 |
| F2-H3 | SRR26987556 |
| F2-H8 | SRR26987555 |
| F2-C1 | SRR26987553 |
| F2-D6 | SRR26987552 |
| F2-H5 | SRR26987551 |

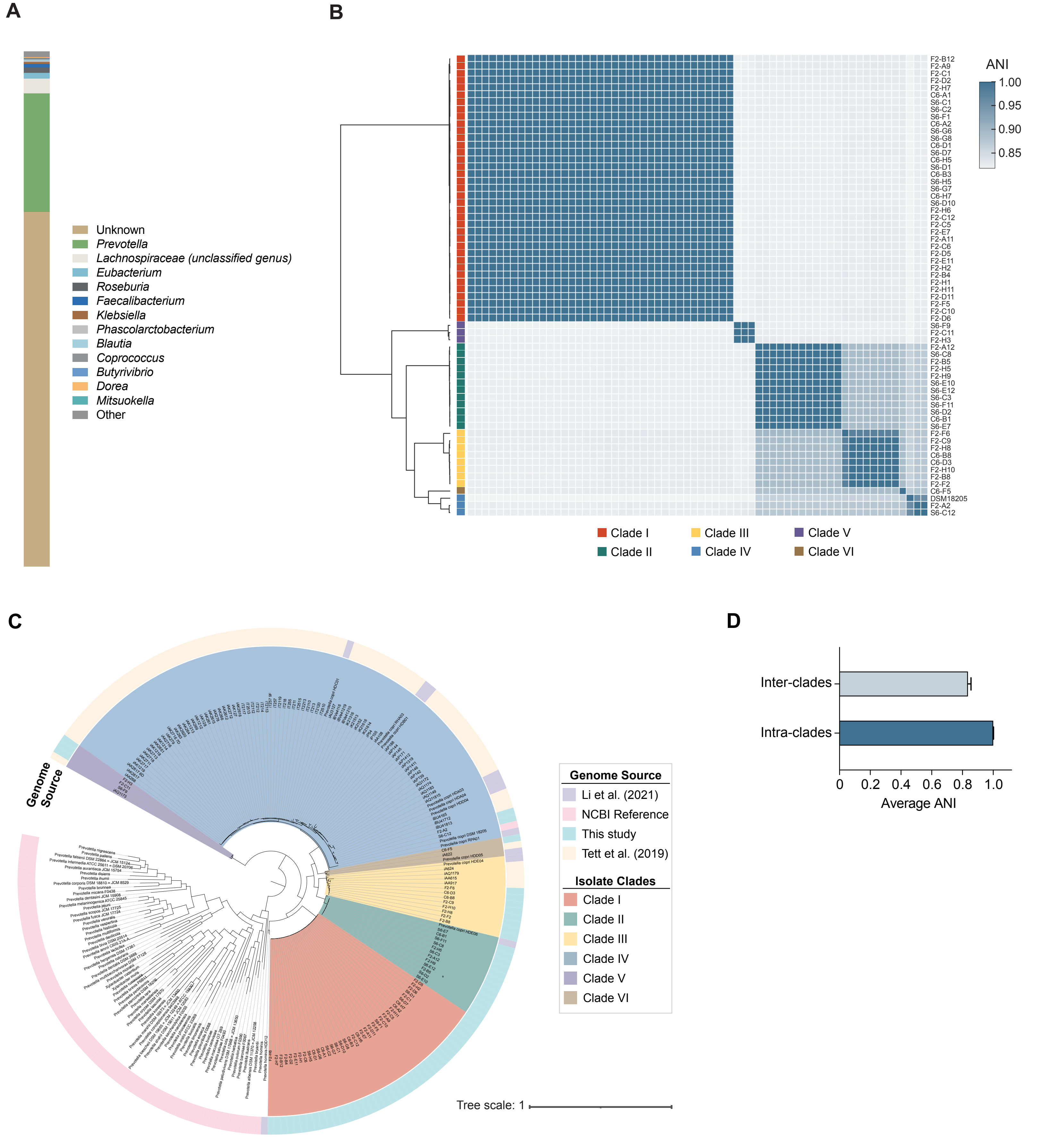


### Supplementary Fig. 1

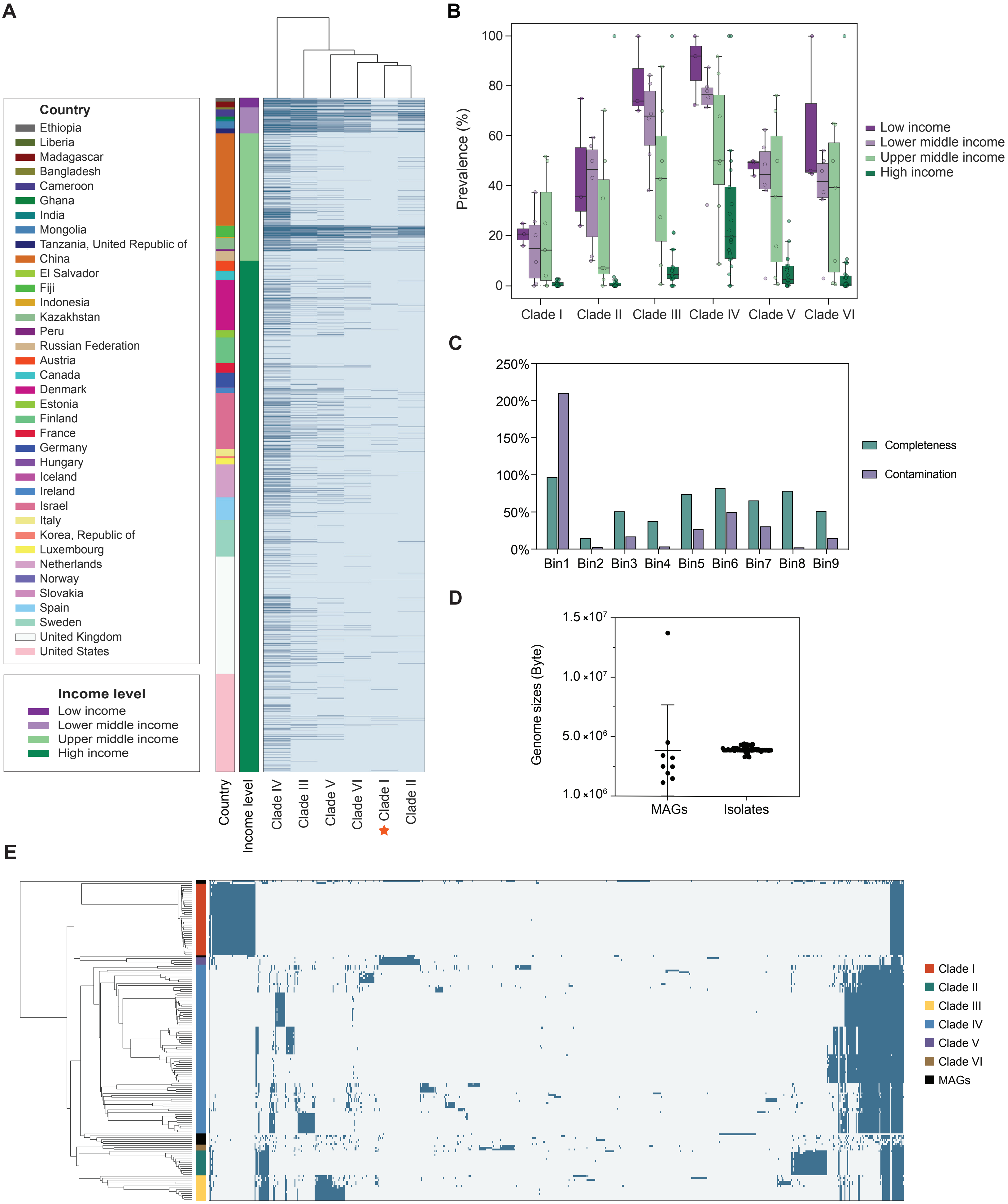


Supplementary Fig. 2

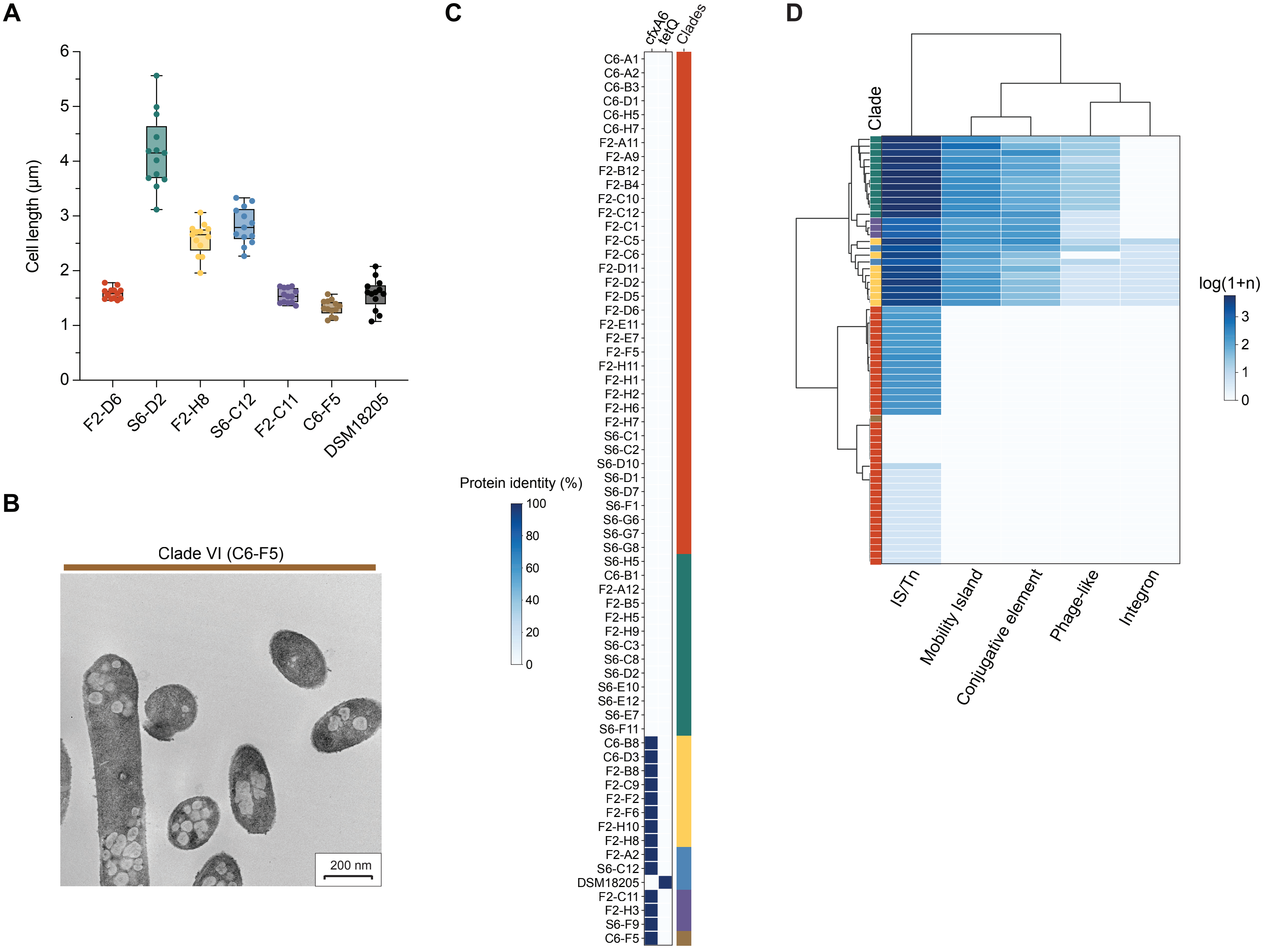


**Supplementary Fig. 3**

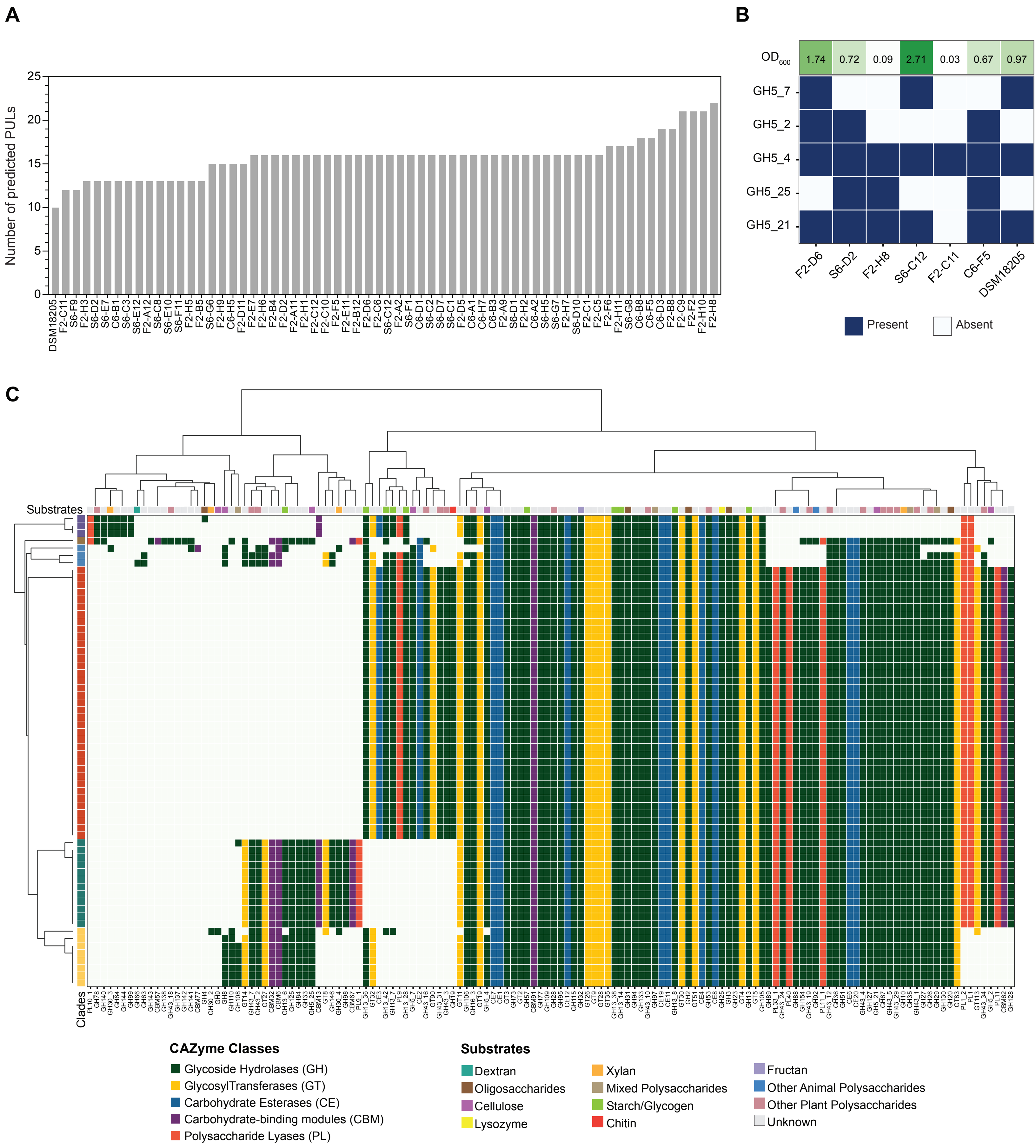


**Supplement Fig. 4**

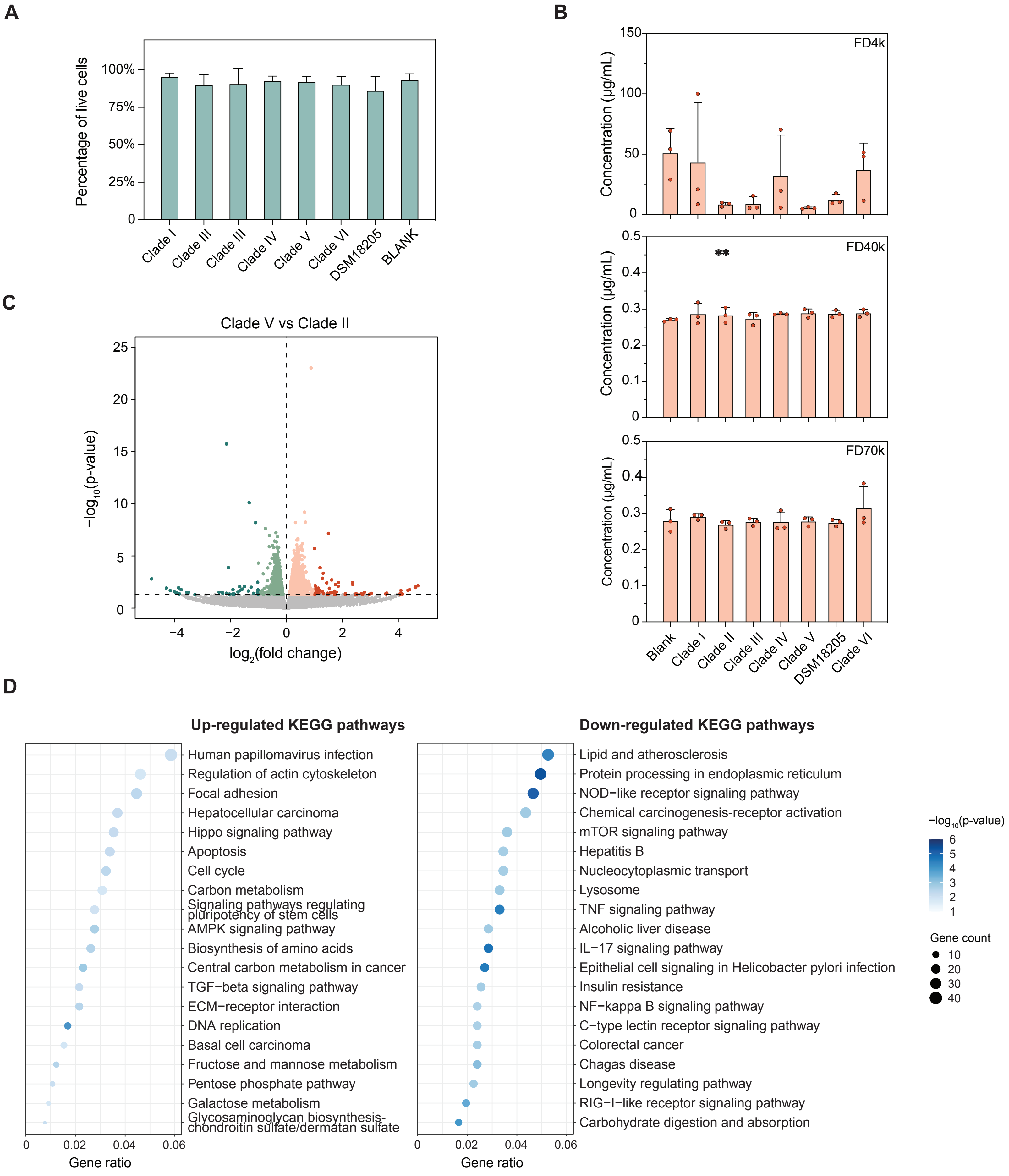


**Supplementary Fig. 5**

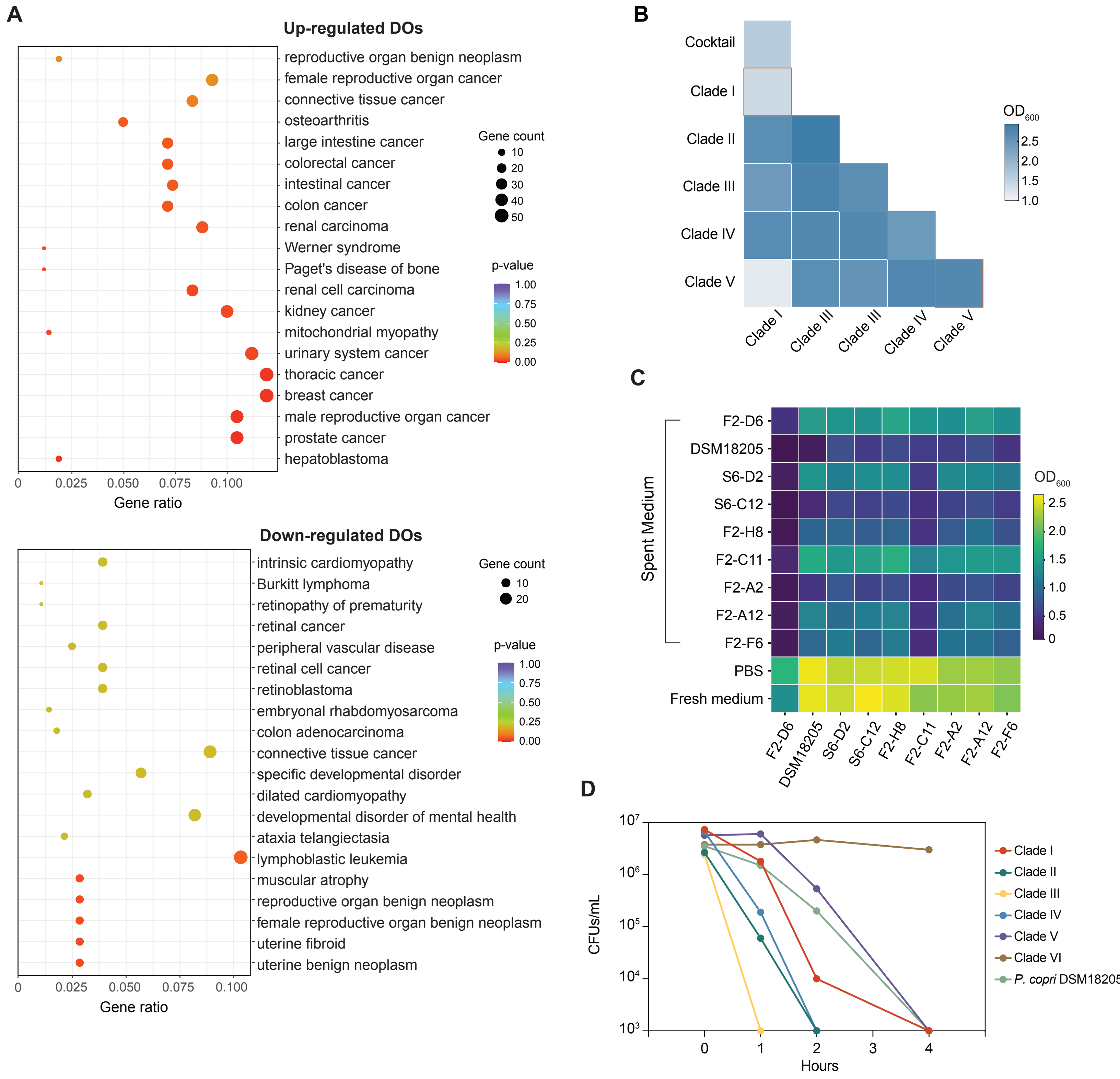


**Supplementary Fig. 6**
